# Conserved Axonal Transcriptome Dynamics Underlie PGE_2_-Induced Sensitisation and Identify *Tnfrsf12a/*Fn14 as a Regulator of Neuronal Excitability in DRG Neurons

**DOI:** 10.1101/2025.09.02.673723

**Authors:** Asta Arendt-Tranholm, Rafael Sebastián Fort, Rebecca Pope, Alex Rathbone, Gareth Hathway, Jose Sotelo-Silveira, Victoria Chapman, Cornelia H. de Moor, Federico Dajas-Bailador

## Abstract

Chronic pain arises when dorsal root ganglion (DRG) neurons become sensitised to noxious inputs, a process driven by inflammatory mediators such as prostaglandin E_2_ (PGE_2_). Local translation of axonal mRNAs is a key regulator of nociceptor plasticity, yet how axonal transcriptome dynamics contribute to inflammatory sensitisation remains unclear. Using compartmentalised culture systems and RNA-sequencing, we defined axonal and somatic transcriptomes in embryonic (E16.5) and adult (W8) DRG neurons and assessed their remodelling after PGE_2_ exposure. We identify a conserved core axonal transcriptome spanning embryonic to adult stages, prominently enriched for ribosomal and mitochondrial functions, consistent with sustained translational and metabolic demands. PGE_2_ elicited compartment-specific reprogramming: pathways related to sensory processing and pain were upregulated in axons but downregulated in somata. Functionally, prolonged axonal PGE_2_ exposure enhanced capsaicin-evoked Ca²⁺ responses and drove retrograde sensitisation of neuronal somata. Integrating transcriptomics with functional assays, we pinpointed *Tnfrsf12a* (Fn14), a cytokine receptor linked to regeneration and neuropathic pain, as a PGE_2_-induced axonal mRNA. Crucially, local axonal knockdown of *Tnfrsf12a* significantly reduced neuronal excitability, providing proof-of-concept that axonally enriched transcripts can be targeted to modulate sensitisation. These findings position conserved axonal transcriptome programmes as drivers of peripheral sensitisation and establish *Tnfrsf12a* as a therapeutic candidate for inflammatory pain.

## Introduction

Chronic pain is a complex clinical condition arising from diverse pathologies, including osteoarthritis, rheumatoid arthritis, migraine, and neuropathy. It manifests as spontaneous pain, exaggerated responses to noxious stimuli (hyperalgesia), or pain evoked by innocuous stimuli (allodynia).

Understanding the molecular mechanisms behind these aberrant responses is difficult due to the interplay of peripheral and central neuronal processes. Sensitisation of nociceptive terminals, originating from dorsal root ganglion (DRG) neurons, is a key driver of the transition from acute to persistent pain and has been a major focus of research (Latremoliere and Woolf 2009; Basbaum *et al*. 2009; Finnerup *et al*. 2021). In this process, the highly polarised cellular structure of DRG neurons constitutes an intrinsic component of this complexity, as it gives rise to distinct subcellular compartments situated in diverse cellular and tissue environments. Indeed, it has become clear that neuronal plasticity and nociception depend on molecular mechanisms capable of integrating and adapting to these specialised functional domains, at both tissue and cellular level. Despite, or partly due to, their intricate anatomical organization *in vivo*, dissociated primary DRG neuronal cultures are widely used as *in vitro* models to investigate neuronal sensitisation, with inflammatory mediators, neuropathic insults, and axonal injury commonly employed to mimic pathological conditions (Chaban 2010; Peeraer *et al*. 2011; Gardiner and Freeman 2016; Chakrabarti *et al*. 2020).

The subcellular RNA localisation and regulated protein expression in dendrites and axons plays a key role in most neuronal plasticity processes, including memory formation, axonal development, regeneration and more recently nociceptor sensitisation (Sahoo *et al*. 2018; Paolo *et al*. 2021; Li *et al*. 2023; Taylor and Nikolaou 2024). The role of localised axonal mRNA translation in sensory axons in the regulation of peripheral nociception has been reported in multiple models of pain (Jiménez-Díaz *et al*. 2008; Price and Géranton 2009; Obara *et al*. 2012; Melemedjian and Khoutorsky 2015; Khoutorsky and Price 2018). Protein translation inhibitors injected at the peripheral terminal but not into the DRG have been shown to reverse nociceptor mediated hyperalgesic priming, implicating different mechanisms in the soma and terminal in the transition to chronic pain (Ferrari *et al*. 2013). Added to this, increasing evidence underscores the role of extracellular cues in regulating local mRNA translation and modulating nociceptor excitability that contributes to the development of hyperalgesia (Price and Inyang 2015; Khoutorsky and Price 2018), with local inhibition of global protein synthesis or direct inhibition of mTOR reducing injury-induced mechanical hypersensitivity and preventing the development of priming (Price *et al*. 2007; Jiménez-Díaz *et al*. 2008).

In this context, experimental models of inflammatory processes are essential for understanding the mechanisms that drive pain and for identifying potential therapeutic targets. Prostaglandin E_2_ (PGE_2_) plays a key role in inflammatory pain linked to infections, autoimmune diseases, and injuries, by activating EP4 receptors on sensory neurons (Southall and Vasko 2001; Clark *et al*. 2008) . As such, it is widely used to model neuronal sensitisation in vitro (Southall and Vasko 2001; Rush and Waxman 2004; Banchet *et al*. 2005; Ma and St-Jacques 2018), and while prostaglandin inhibition can reduce inflammatory pain, chronic NSAID use carries serious side effects, underscoring the need for alternative therapeutic targets (Ricciotti and FitzGerald 2011).

In this study, we used compartmentalised culture systems to investigate the molecular and subcellular mechanisms underlying inflammatory pain. By analysing both embryonic and adult DRG neurons in specialised microchambers, we uncovered localised changes in the axonal mRNA transcriptome following PGE_2_-induced nociceptive sensitisation. Notably, our combined analytical and experimental approach identified a role for axonal FN14, encoded by *Tnfrsf12a*, in modulating DRG neuron excitability. These findings underscore the importance of subcellular transcriptomics in dissecting peripheral pain mechanisms and establish a novel pipeline for identifying axonally regulated targets in sensory neuron sensitisation.

## Materials and Methods

### Animals

C57BL/6J mice (Charles River) were bred and group-housed at the University of Nottingham, UK, in accordance with the Animals (Scientific Procedures) Act 1986/2012 or at the Animal Research Facility in accordance with Institutional Animal Care and Use Committee at the University of Texas at Dallas. Embryonic day 16.5 (E16) mice, extracted from pregnant dams, and 8-week-old mice (W8) were kept and sacrificed under schedule 1 procedures conducted according to the relevant guidelines for age and species.

### DRG culture

Spinal cords from E16.5- and 8-week-old mice were gently removed from each pup, using forceps, to provide a clear view of the DRGs, and these were individually extracted from the intervertebral foramen using forceps and placed into Leibovitz’s L-15 (1X, Gibco).

DRG explants were transferred to 0.025% trypsin w/v (T9201, Sigma) in Dulbecco’s Phosphate Buffered Saline (MgCl_2_- and CaCl_2_-free PBS, D8537 Sigma) and incubated for 20-minutes at 37°C.

Explants were then transferred into filter-sterilised 0.2% (w/v) collagenase type II solution (17101- 015, Gibco), in MgCl_2_- and CaCl_2_-free PBS, and incubated for 20-minutes at 37°C. DRG explants were removed from collagenase and added to 1 mL resuspension media: D6546 DMEM, heat-inactivated FBS (10%), glutaMAX (1%, 1X Gibco) and Penicillin-Streptomycin (1%, Sigma).

Gentle mechanical disruption was used to dissociate aggregated cells and resuspend cells in a homogeneous solution. After a 5-minute centrifugation at 4000 rpm, the supernatant was discarded and the cell pellet was gently resuspended in the required volume of complete media: D6546 DMEM, glutaMAX (1%, 1X Gibco), B27 supplement (2%, ThermoFisher Scientific), Penicillin/Streptomycin (1%, Sigma), Nerve Growth Factor 2.5S (NGF, 50 ng/mL, ThermoFisher Scientific), Glial-Derived Neurotrophic Factor (GDNF, 50 ng/mL, Sigma) and Aphidicolin (APH, 50 ng/mL, Sigma) for seeding. Complete media was supplemented with NGF and GDNF to promote neuronal proliferation and axonal outgrowth, and the DNA polymerase inhibitor APH to reduce the proliferation of non-neuronal cells.

Nunc dishes (Nunclon™ Delta 35x10 mm, ThermoFisher Scientific) were coated with 0.02 mg/mL poly-L-lysine (PLL, Sigma P1524) for dissociated cultures and 0.1 mg/mL PLL for compartmentalised microfluidic cultures (Xona Microfluidics, SND150), the day before dissection. Dishes were incubated at 37°C for 1-2 hours, washed twice with dH_2_O, and airdried overnight in a class II cabinet. Following dissection and prior to seeding, custom polydimethylsiloxane (PDMS) rings (11mm diameter) or compartmentalised microfluidic chambers were applied to nunc dishes. Microfluidic channels and the centre of PDMS rings were coated with 20 µg/mL laminin (in DMEM, D6456 Sigma), and incubated at 37°C for at least an hour before removal.

100 µL complete media was added to the axonal-enriched chamber compartment before seeding. Dissociated neurons from 15 DRGs were added to each chamber, suspended in 20 µL complete media, by pipetting 10 µL complete media cell solution into each end of the soma-enriched channel. Chambers were incubated at 37°C for at least two hours before an additional 200 µL of complete media was added to the soma compartment. The following day, 50 µL of media was transferred from one somal well to the other to remove any unattached cells. Media was changed every 2-3 days.

The protocol for seeding and culturing in porous membrane chambers was adapted from previous experimental protocols (Unsain *et al*. 2014). Porous membrane chambers with 1µm pores (Corning Life Sciences, 353102) were coated with 0.1 mg/mL PLL in dH_2_O for 1 hour and dried overnight. Laminin-coating of 20 µg/ml in DMEM was carried out 1 hour prior to cell seeding. 25 dissociated DRG-explants (6.39x105 cells/ml) were seeded above membrane in 100μl complete media. DRG-cells were allowed to grow for up to 7 days to facilitate the development of an axonal network below the membrane, with media changes every 2-3 days. DRG-cells were grown in incubators at stable conditions of 37°C and 5% CO2.

RNA extraction from E16.5 cells in porous membrane chambers was carried out according to an experimental protocol from (Garcez *et al*. 2016) with TRIzol™ Reagent (Invitrogen 15596018) and chloroform. Each side of the membrane was scraped gently to promote detachment and a guanidine thiocyanate wash (GTC solution, Sigma Aldrich 50983, in Sodium Acetate and Isopropanol) was performed to minimize protein and RNase contamination. RNA extraction from W8 cells in porous membrane chambers was performed with the TRIzol-based RNA Miniprep Plus (Zymo Research) at the University of Texas at Dallas according to manufacturer instructions. DNase I treatment was performed on all RNA extractions prior to RNA sequencing. Three independent biological replicates were included for sequencing for both E16.5 and W8 cell-cultures.

### PGE_2_ Treatment

Prostaglandin E_2_ (PGE_2_, Cayman Chemical) or the stabilized derivative 16,16-dimethyl prostaglandin E_2_ (16,16-PGE_2_, Tocris) in dimethyl sulfoxide (DMSO, D4540 Sigma) was purged with nitrogen and a 10 mM stock solution established through dilution in MgCl_2_- and CaCl_2_-free PBS. A final working concentration of 10 μM was obtained fresh by diluting in DRG complete media. E16.5 cultures were exposed to PGE_2_ with replenishment every 2H, or 16,16-PGE_2_ with replenishment every 12 hours, to maintain concentration of 10 μM for 24H prior to RNA extraction. To accommodate logistical constraints, W8 cultures were limited to 12-hour PGE_2_ exposure prior to RNA extraction.

Dissociated neurones were cultured for 7 days before whole-cell incubation with 10 μM PGE_2_ 24- hours. In compartmentalised microfluidic cultures, DRGs were cultured for 6 days *in vitro* to ensure extensive axon crossing through the microgrooves to the axon-enriched compartment. 17-hours prior to calcium imaging, media was removed from the axonal compartment of the chamber and replaced with complete media containing 10 μM stabilised 16,16-dimethyl PGE_2_, or vehicle control, to simulate a peripheral insult. Cultures were returned to the incubator until Ca^2+^ imaging was performed the following day.

### Tnfrsf12a siRNA Treatment

A SMARTPool of four Accell siRNAs (Horizon Biosciences) targeting murine *Tnfrsf12a* mRNA was used to evaluate gene knockdown. A non-targeting siRNA pool (Accell Non-Targeting Control Pool, Horizon Biosciences) served as a control to account for off-target effects, while a media change-only group (untreated control) was included to account for baseline gene expression. Both siRNA pools were reconstituted in HyClone water to a 100 μM stock, mixed on a roller mixer for 30 minutes at room temperature, aliquoted, and stored at -20 °C. Prior to treatment, siRNAs were diluted in DRG complete media to a final concentration of 1 μM. Primary DRG neurons, dissociated from 20 ganglia and cultured in 6-well plates, were treated on DIV5 with either *Tnfrsf12a* siRNA or non-targeting control siRNA. RNA was extracted at 24 and 48 hours post-treatment (DIV6 and DIV7) using TRIzol® Reagent (Invitrogen), as described previously. RNA quality and quantity were assessed via NanoDrop spectrophotometry; samples with a 260/280 absorbance ratio near 2.0 were considered acceptable.

cDNA synthesis was performed using SuperScript™ IV Reverse Transcriptase (Invitrogen) with oligo(dT)20 primers, and reactions were run on an Eppendorf Mastercycler. Quantitative PCR (qPCR) was carried out using PowerUp™ SYBR™ Green Master Mix (Applied Biosystems) on a QuantStudio 5 system. Expression of *Tnfrsf12a* was quantified relative to the geometric mean of two validated housekeeping genes, *Gapdh* and *Ube2*, using the comparative Ct method (2^ΔΔCt^) (Schmittgen and Livak 2008).

### Ca^2+^ Imaging

Ca^2+^ transients within the cell soma were quantified as a measure of neuronal excitability, following stimulation. DRGs in compartmentalised microfluidic devices or dissociated cultures were incubated in 1 μM Fluo-5 AM (Sigma-Aldrich, diluted in Pluronic F127 20% solution in DMSO) for 30-minutes at RT and then de-esterified in 1X imaging buffer (135 mM NaCl, 3 mM KCl, 10 mM HEPES, 15 mM glucose, 1 mM MgSO_4_, 2 mM CaCl_2_, in PBS) for a further 30-minutes at RT. All stimuli were reconstituted in 1X imaging buffer.

Nunc dishes were directly mounted onto the microscope stage and an image acquisition area selected; for compartmentalised microfluidic chambers, an area within the soma-enriched compartment, close to the entrance of the microgrooves, was chosen and for dissociated cultures, a field of view near to the centre of the dish.

Fluorescent images were acquired at 488 nm excitation every 500 ms, with a 100 ms exposure period, using Channel LED Green 470. An Olympus IX70 inverted tissue culture microscope was attached to a CoolLED pE excitation system, PE-2 collimator and an ORCA-R2 Hamamatsu Camera Controller C10600. Image recording was conducted through multi-acquisition in ImageJ, with 1200 frames recorded for 10 minutes in dissociated cultures and 1350 frames recorded for 11 minutes and 15 seconds for compartmentalised cultures (one frame every 500 ms to minimise photobleaching). Baseline measurements were taken over the first 15-seconds of recording, before the first stimulation at 30 seconds. Stimuli were gently added directly to cells in dissociated dishes, or to the appropriate channel compartment of compartmentalised devices, after 30 seconds of recording. In microfluidic cultures, a second stimulation of 25 mM KCl was added to the soma side.

### Data Analysis

#### RNA-sequencing

DNase treatment was carried out; RNA purity and integrity were assessed using an Agilent 2100 Bioanalyzer System using the RNA 6000 Nano assay (Agilent Technologies). All samples had a RIN value above 7. For embryonic DRG neurons, library preparation of rRNA depleted RNAs was prepared for Illumina HiSeq paired-end 150bp sequencing by GeneWiz (Azenta Life Sciences). For adult DRG neurons, library preparation of polyA selected mRNAs was prepared for Illumina HiSeq single-end 75bp sequencing by the University of Texas at Dallas Genome Core Next Seq 500. Quality checks of the RNA-seq data were performed using FastQC. Filtered high quality reads from samples (embryonic DRG neuron Ave. 28,442,101 ± SD 12,577,154 reads and Adult DRG neurons Ave.

32,947,254 ± SD 5,322,518 reads, **Supplementary Table 1**) were mapped to the GENCODE GRCm38 genome using the GENCODE vM25 reference annotation through the STAR mapping tool-package (embryonic DRG neuron overall genome mapping: Ave. 85.1 ± SD 11.2, and Adult DRG neurons Overall genome mapping: Ave. 98.4 ± SD 0.2, **Supplementary Table 1**). HTSeq was subsequently used to obtain counts tables for further analysis. Filtering was conducted to only consider protein-coding, non-mitochondrial genes. Normalisation and Differential Expression Genes (DEGs) analysis were performed using the SARTools pipeline in RStudio, selecting the edgeR algorithm, Trimmed Mean of the M-values (TMM) normalisation and CPM (Counts Per Million) ≥ 1 as the cut-off (for edgeR analysis results of E16.5 embryo dataset see **Supplementary Table 2** and **Supplementary Figure 1**; and **Supplementary Table 3** and **Supplementary Figure 2** for W8 adults). On average, 85% of reads from embryonic DRG neurons and 98% of reads from adult DRG neurons were mapped to the genome (see Methods, **Supplementary Table 1-2** and **Supplementary Figures 1-2**). Although both datasets exhibit a high percentage of global mapping, there was a distinction in the average mapping rates between embryonic and adult DRG samples likely due to differences in library construction strategies, specifically rRNA depletion versus poly-A selection. Since no differentially expressed genes were obtained after multiple testing correction (adjusted p-value), we used the nominal p-value (unadjusted) as a reference to visualise global transcriptional changes. Volcano plots were generated only to display genes with ≥1.5-fold change and p < 0.05 for PGE_2 vs_ Control; however, these genes were not used for biological interpretation. Instead, gene set enrichment analysis (GSEA) was applied, which is independent of p-value thresholds and considers the entire ranked list of genes to identify functionally enriched pathways (Wu *et al*. 2021). The raw FASTQ data sets supporting the results of this article is available at the Sequence Read Archive repository (https://www.ncbi.nlm.nih.gov/sra), BioProject ID: PRJNA1337668.

#### Gene Set Enrichment Analysis

Gene Set Enrichment Analysis (GSEA) was used to assess the enrichment of entire biological pathways, rather than relying solely on discrete differentially expressed genes (DEGs). By reducing dependence on arbitrary significance thresholds, GSEA captures subtle yet biologically meaningful transcriptional patterns, allowing for a more comprehensive interpretation of the dataset (Subramanian *et al*. 2005; Wu *et al*. 2021). GSEA was performed using gseGO (R package clusterProfiler v4.8.2) to explore the Gene Ontology (GO) terms (Biological Process, Molecular Function and Cellular Component) using the complete list of genes (ranked based on fold change of expression). The terms, pathways, or networks with a significance threshold of adjusted p-value < 0.05 were selected. Figures were designed using R and all analyses were performed in R software (version 4.3) using libraries: clusterProfiler, enrichplot, ggplot2, heatmap.2 (clustering distance measured by Euclidean and Ward clustering algorithms).

#### Ca^2+^ Imaging

Fluorescence emitting cells were manually selected using the oval tool on FIJI ImageJ but were excluded from analyses if the response did not surpass five standard deviations of baseline fluorescence, measured over 15-seconds prior to stimulation. Each biological replicate contained at least two replicates per treatment with at least three responsive cells per technical replicate. Data were processed on Excel. All statistical analyses were performed, and all graphs produced, on GraphPad PRISM 10. Student’s t-tests were performed on normally distributed data. A p-value <0.05 was considered significant (*) and a p-value <0.01 considered very significant. Ca^2+^ traces depict the ΔF/F_0_ averaged across three biological replicates (each containing at least 3 technical replicates).

Peak values refer to the maximum ΔF/F_0_ of individual cells within technical replicates, averaged across the three biological replicates.

## Results

### PGE_2_ driven increase in DRG neuron excitability

To begin to investigate the cellular and molecular mechanisms underlying neuronal excitability, dissociated DRG neurons from E16.5 mice were cultured for seven days on 35 mm dishes, and their responses to capsaicin (200 nM) was assessed via fluorescence imaging of Ca²⁺ transients in the soma/cell body. A population of small-diameter DRG neurons (100–300 μm²) responded to capsaicin, consistent with their classification as nociceptors and confirming their correspondence to small C-fiber nociceptors. To establish a sensitisation model, dissociated DRG neurons (**Fig 1A**) were incubated with PGE_2_ (10 μM, 24 h), prior to stimulation with capsaicin (200 nM). DRG neurons pre-incubated with PGE_2_ showed an increase in Ca²⁺ responses to capsaicin stimulation (**Fig 1D**), evident in the significantly higher peak fluorescence intensity (**Fig 1G**), and as a marked increase in overall response magnitude, quantified as area under the curve (**Fig 1J**). Confirming the effects of PGE_2_, we also detected a significant increase in *Ngf* mRNA levels in the DRG neurons relative to the housekeeping gene *Hprt* (2DDCt, *p=0.0232*, n=5), corroborating the activation of neuroplastic pain pathways in our in vitro model.

**Figure 1.**
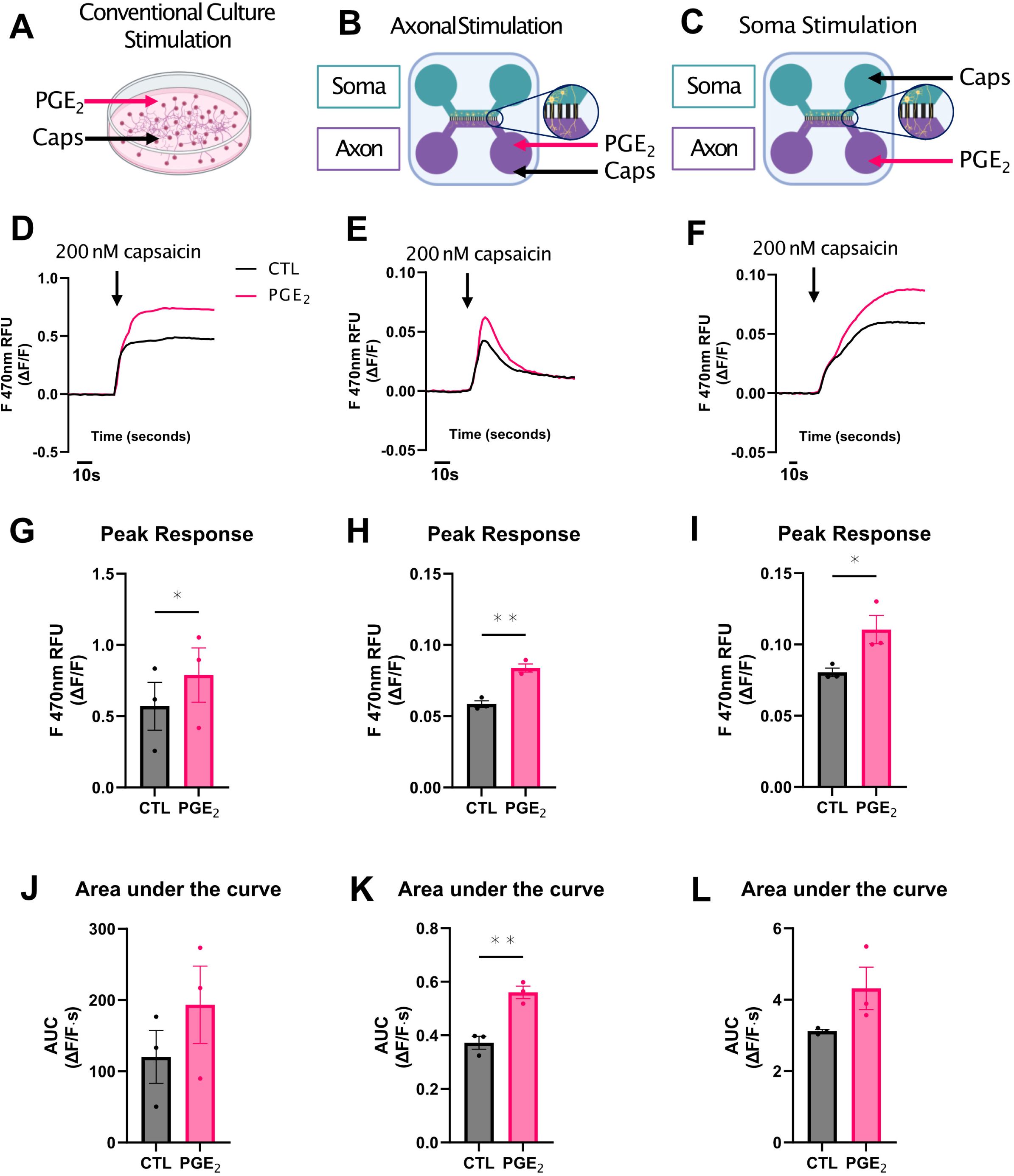
PGE_2_ enhances capsaicin-evoked excitability of DRG sensory neurons as shown by increased Ca^2+^ transients in the soma. **A-C**. Schematics of the experimental set-ups. **A.** Dissociated DRG sensory neurons were cultured for 7 days and incubated with 10 μM PGE_2_ for 24 hours. Ca^2+^ imaging was then performed following stimulation with 200 nM capsaicin. **B-C.** In compartmentalised microfluidic cultures, PGE_2_ was applied selectively to the axonal compartment for 17 hours. During calcium imaging, capsaicin stimulation was applied to either the same axon (B) or soma compartment (C). **D-F.** Average ΔF/F_0_ Ca^2+^ traces measured in the soma in response to capsaicin stimulation. **D.** PGE_2_-treated neurons in dissociated cultures showed an enhanced Ca^2+^ response. **E.** In the microfluidic set-up, axonal stimulation produced a heightened response in PGE_2_-treated neurons, with a characteristically narrower temporal profile. **F.** Direct somatic stimulation with capsaicin also led to an enhanced Ca^2+^ response in the PGE_2_ group. The trace shows a more prolonged signal, typical of direct cell body stimulation. **G-I.** Quantification of calcium transients (peak response, ΔF/F_0_) revealed significant increases in the PGE_2_ group across all stimulation types: dissociated cultures (G), axon stimulation in microfluidic chambers (H), and soma stimulation (I) (*t*-test, *p* < 0.005). **J-L.** Analysis of area under the curve showed a marked increase in PGE_2_ treated neurons, with statistically significant changes only following axonal stimulation (**K**, *t*-test; *p* < 0.05), but not in dissociated cultures **(J)** or following soma stimulation **(L)**. For all panels, *N* = 3 biological replicates, with ≥ 2 technical replicates. Data are presented as mean ± SEM. *p* < 0.05 was considered statistically significant.

To assess whether PGE_2_ can also drive sensitisation at axon terminals, DRG neurons were cultured in compartmentalised microfluidic chambers, where by day six they extended substantial axonal projections (**Fig 1B**). Fluidic isolation enabled selective stimulation of axons, revealing that capsaicin (200 nM) applied only to axon terminals elicited somatic Ca²⁺ transients of shorter duration than whole-cell stimulation (**Fig 1E**). Axonal exposure to prolonged PGE_2_ (10 μM, 17 h) significantly enhanced these axonal stimulations to capsaicin (**Fig 1E**), triggering significantly larger somatic Ca²⁺ transients compared to naïve axons (**Fig 1H** & **1K**), both in peak and duration of response. Notably, axonal PGE_2_ followed instead by somatic capsaicin stimulation (**Fig 1C** & **1F**) also heightened somatic responses (**Fig 1I** & **1L**). These findings identify an axon-driven mechanism of neuronal sensitisation and emphasise the need to dissect local transcriptomic programs that coordinate signalling between axons and somata.

### Characterisation of local transcriptomes from DRG neurons in culture

To investigate potential local changes in subcellular transcriptomes, both at somatic and axonal level, we implemented a DRG neuron culture using modified Boyden chambers with a porous membrane that allows the growth of axonal projections separately from the DRG neuronal cell bodies. To confirm the nature and cellular origin of the RNAs being analysed, we investigated the expression of different cell type markers within our RNA-seq data (**Supplementary Fig 3**). These included 4 markers for Schwann cells, 12 markers for satellite glial cells and 11 markers for DRG neurons, as based on previous studies characterising the cell-subtypes of the dorsal root ganglia (Zeisel *et al*. 2018; Ray *et al*. 2018; Wangzhou *et al*. 2020; Liang *et al*. 2020). As shown in **Supplementary Fig 3A**, the soma transcriptomes of E16.5 cultures exhibit high expression of most DRG neuron markers, while markers for Schwann cells and satellite glial cells remain comparatively low. This confirms that our culture system is primarily composed of DRG neurons, ensuring a representative model for studying their transcriptomic profiles.

Unlike this characteristic expression of DRG markers found in the soma, RNA expression patterns in the axon compartment showed a more heterogeneous population of neuronal/non-neuronal markers (**Supplementary Fig 3A**). Although this could suggest potential ’contamination’ from other cell types, the relative absence of such markers in the somatic transcriptome makes this unlikely.

Alternatively, the data might reflect a diversity in axonal transcriptomes which does not strictly conform with a canonical cell-type-specific signature and instead reflects the distinct functional requirements of the axonal compartment. Based on this, we further analysed the expression of a selected group of genes previously identified as enriched in neurite-specific transcriptome studies (Kügelgen and Chekulaeva 2020). Crucially, we observed elevated expression of all selected neurite marker genes in our axonal transcriptomes relative to the soma counterpart (**Fig 2A**), supporting the view that these transcriptomes reliably represent the axonal subcellular domain. Taken together with previously published axonal transcriptomes, our data also points to a conserved molecular identity that is shared across neuronal types.

**Figure 2.**
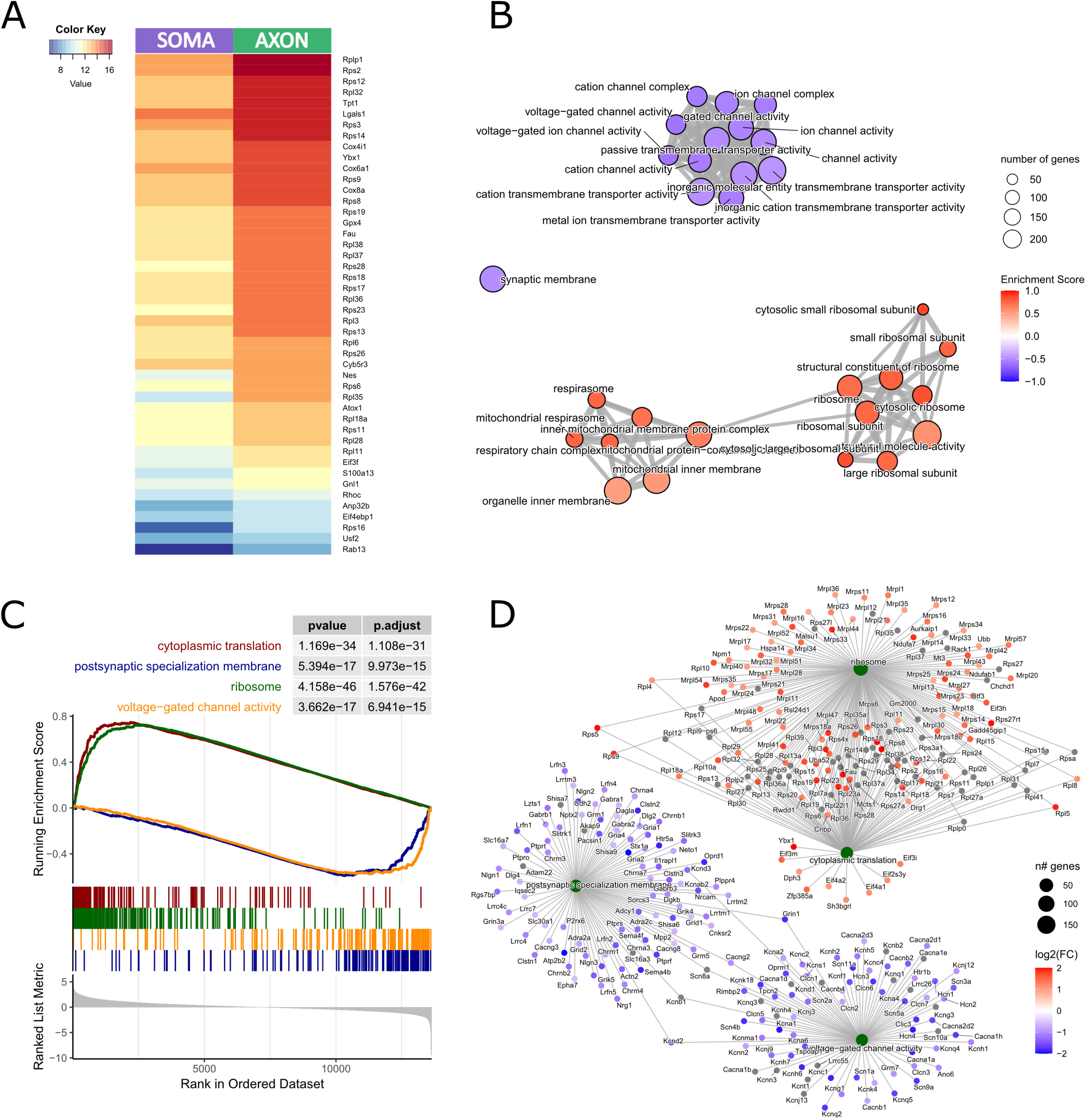
Analysis of gene expression in the Axon and Soma compartments of E16 DRG neurons. **A.** Heatmap of neurite marker genes (*Kugelgen & Chekulaeva, 2020*), ordered from most to least abundant in the axon compared to the soma. **B.** Emmaplot of the top 30 significantly enriched GO Biological Process terms from the GSEA analysis comparing Axon vs Soma. Circle size represents the number of genes associated with each term, while the color scale indicates whether the term is significantly enriched in the axon (red) or depleted in the axon (blue) relative to the soma. **C.** GSEA plot of selected terms. The x-axis represents the ranked list of all genes; the y-axis (upper panel) shows the Enrichment Score, and the y-axis (bottom panel) displays the ranking metric value. **D.** Cnetplot of the selected terms from panel C, showing the genes associated with each enriched term. Circle color represents the log2FC value from the axon vs soma comparison.

To probe transcriptomic differences between axonal and somatic compartments, we performed Gene Set Enrichment Analysis (GSEA) using Gene Ontology (GO) terms spanning Biological Process, Molecular Function, and Cellular Component. Genes were ranked by fold-change between axons and soma, and enrichment scores identified overrepresented functional categories. These were visualised with an Enrichment Map, enabling assessment of physical and functional associations among pathways based on shared genes. Axonal transcripts were significantly enriched for GO terms related to protein synthesis and mitochondrial function, including *structural constituent of ribosome*, *ribosomal subunit*, *cytoplasmic translation*, *translation at pre-synapse*, *mitochondrial inner membrane*, *respirasome*, and *respiratory chain complex* (**Fig 2B**). To highlight some of these trends, we analysed gene set distributions within selected pathways: transcripts linked to *ribosome* and *cytoplasmic translation* clustered among the most enriched axonal genes, whereas those associated with *post-synaptic specialised membrane* and *voltage-gated channel activity* were concentrated at the bottom of the ranked list, reflecting relative depletion in axons (**Fig 2C**).

When inspecting in more detail the identity of the genes included in these selected ontology terms, we could observe a high enrichment of mRNAs coding for ribosomal proteins (RPs) in the axonal compartment (**Fig 2D**) compared to the soma, an experimental finding that is consistent with previous studies showing RPs among the most upregulated/enriched in the axonal transcriptome (Deglincerti and Jaffrey 2012; Shigeoka *et al*. 2019). Remarkably, Rps4x, whose axonal synthesis was shown to be crucial for axon development *in vivo* (Shigeoka *et al*. 2019) was amongst the most enriched RP in our axonal transcriptomes from E16 derived DRG neurons (**Fig 2D**), reinforcing the potential importance of ribosome remodeling in sensory axons. Another key RNA/DNA binding protein, YBX1, with known roles in the axon and defined as neurite-enriched in (Kügelgen and Chekulaeva 2020), was among those within the *cytoplasmic translation* ontology term, highlighting a potentially important role of these proteins in axonal development and function.

### Conservation of Axonal Transcriptome Signatures from Embryonic to Adult DRG Neurons

An expanding body of research continues to underscore the critical role of axonal mRNA transcripts in supporting axonal function within the highly polarised architecture of neurons. However, most experimental investigations to date have concentrated on developmentally active, embryonically derived neurons, such as the E16.5 neurons examined so far in this study, leaving the molecular dynamics of mature neuronal axons comparatively underexplored. To determine whether the transcriptomic profiles observed in E16.5 axons are also present at later stages, we analysed the axonal transcriptomes in DRG primary neurons from 8-week-old adult mice (W8). As shown in **Suppl Fig 3B**, most of the DRG neuron markers identified in our E16 studies are also highly expressed in the soma transcriptomes of W8 cultures, while markers for Schwann cells and satellite glial cells are comparatively low. Also consistent with our findings in E16.5-derived DRG neurons, W8 axonal transcriptomes exhibit comparatively high expression of nearly all neurite marker genes (**Fig 3A**), and show prominent enrichment for *translation*, *ribosome-associated* pathways, and *mitochondrial* functions (**Fig 3B**-**D**), while transcripts related to *postsynaptic specialization* and *voltage-gated channel activity* are comparatively less abundant. These findings reveal a conserved molecular signature that likely supports essential axonal functions such as local protein synthesis and energy production. The overlap in basal transcriptomic identities between early and late-stage DRG neurons suggests sustained functional requirements across development and into adulthood, while also raising the possibility that, under *in vitro* conditions, both embryonic and adult neurons may need to express and recapitulate key developmental programs that support axonal integrity and adaptability.

**Figure 3.**
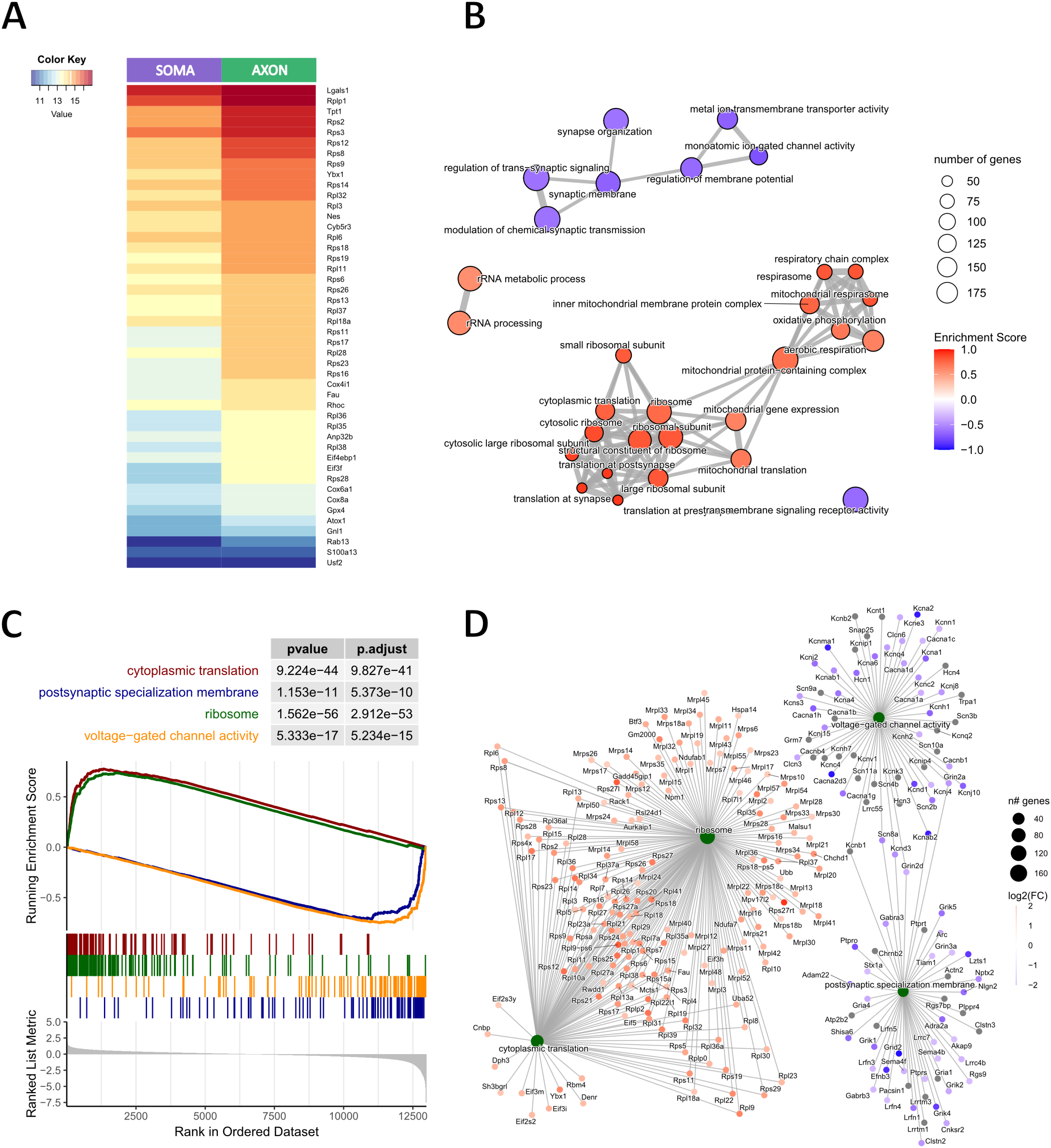
Analysis of gene expression in the Axon and Soma compartments of adult DRG neurons. **A.** Heatmap of neurite marker genes (*Kugelgen & Chekulaeva, 2020*), ordered from most to least abundant in the axon compared to the soma. **B.** Emmaplot of the top 30 significantly enriched GO Biological Process terms from the GSEA analysis comparing Axon vs Soma. Circle size represents the number of genes associated with each term, while the color scale indicates whether the term is significantly enriched in the axon (red) or depleted in the axon (blue) relative to the soma. **C.** GSEA plot of selected terms. The x-axis represents the ranked list of all genes; the y-axis (upper panel) shows the Enrichment Score, and the y-axis (bottom panel) displays the ranking metric value. **D.** Cnetplot of the selected terms from panel C, showing the genes associated with each enriched term. Circle color represents the log2FC value from the axon vs soma comparison.

### PGE_2_ driven changes in local transcriptomes

Considering the differences in transcript enrichment between axons and somas, and the increasing recognition of the importance of subcellular transcriptomes in the dynamic regulation of neuronal function, we investigated the effects of PGE_2_ exposure on the axonal and somal transcriptomes. We found that those axonal markers previously identified in DRG neurons (**Fig 2**) exhibited compartment-specific changes in response to PGE_2_ stimulation, with their expression decreasing in the axonal compartment and increasing in the soma (attenuation of their axonal profile), compared to controls, in both E16.5 and W8 DRG neurons (**Figs 4A and 5A**). These compartment-specific shifts suggest a redistribution or altered trafficking of axonal transcripts in response to PGE_2_ stimulation, potentially reflecting dynamic remodelling of the local transcriptome to support context-dependent functional changes, such as sensitisation or injury-like responses, in DRG neurons. Remarkably, in both E16.5 and W8 DRG neurons, hierarchical clustering of transcriptomic profiles first segregates samples by subcellular compartment, axon versus soma, prior to distinguishing between PGE_2_- treated and control conditions. This highlights the compartmentalised organisation of neuronal transcriptomes and reinforces the existence of a distinct axonal mRNA signature (**Fig 4-5**)

**Figure 4.**
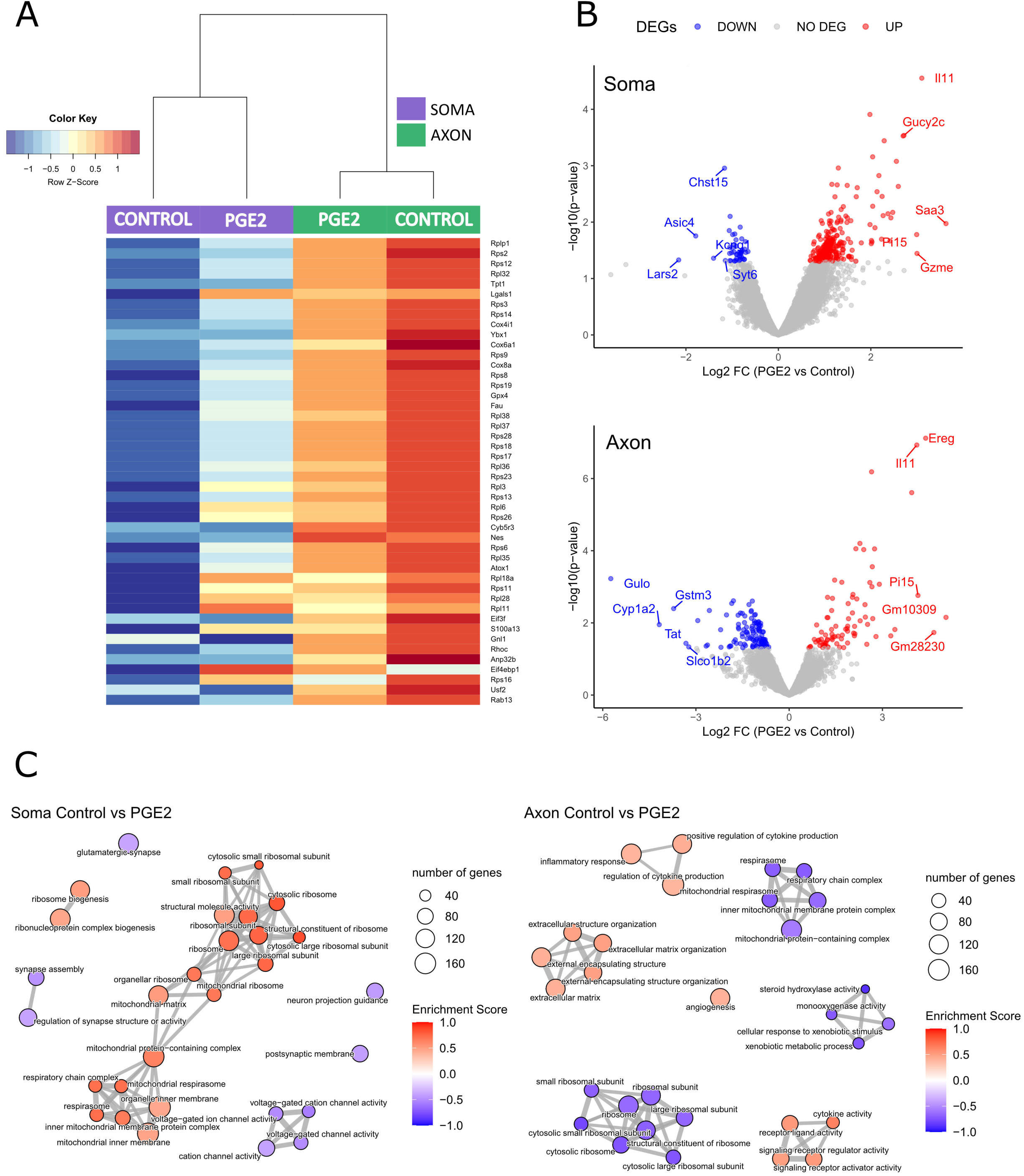
Analysis of gene expression after PGE_2_ exposure on the axonal and somal transcriptomes of E16 DRG neurons. **A.** Heatmap of neurite marker genes (Kugelgen & Chekulaeva, 2020) comparing soma (violet) and axon (green) compartments after PGE_2_ exposure. **B.** Volcano plots of the PGE_2_ vs control comparisons for the Soma and Axon compartments. Genes with a log2FC greater than |0.58| and a p-value lower than 0.05 are highlighted in color; the names of the top 5 upregulated and downregulated genes are indicated. **C** and **D.** Emmaplots of the top 30 significantly enriched GO Biological Process terms from the GSEA analysis comparing PGE_2_ vs Control for Soma (**C**) and Axon (**D**). Circle size represents the number of genes associated with each term, while the color scale indicates whether the term is significantly enriched (red) or depleted (blue).

**Figure 5.**
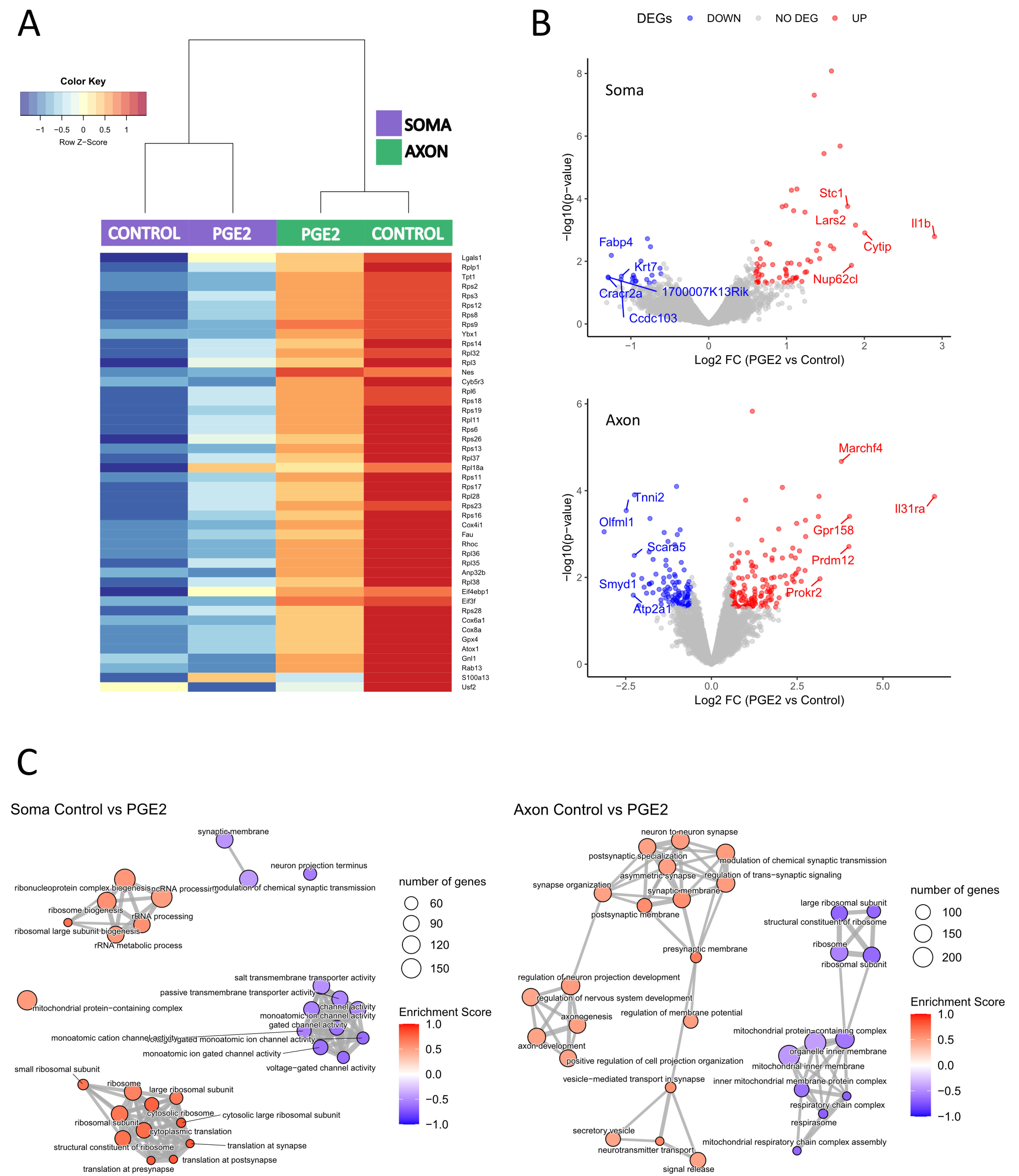
Analysis of gene expression of PGE_2_ exposure on the axonal and somal transcriptomes of adult DRG neurons. **A.** Heatmap of neurite marker genes (Kugelgen & Chekulaeva, 2020) comparing soma (violet) and axon (green) compartments after PGE_2_ exposure. **B.** Volcano plots of the PGE2 vs control comparisons for the Soma and Axon compartments. Genes with a log2FC greater than |0.58| and a p-value lower than 0.05 are highlighted in color; the names of the top 5 upregulated and downregulated genes are indicated. **C** and **D.** Emmaplots of the top 30 significantly enriched GO Biological Process terms from the GSEA analysis comparing PGE_2_ vs Control for Soma (**C**) and Axon (**D**). Circle size represents the number of genes associated with each term, while the color scale indicates whether the term is significantly enriched (red) or depleted (blue).

Beyond the changes observed in this signature of axon-specific markers, E16.5 neurons exhibited marked transcriptomic changes, with numerous genes upregulated or downregulated in response to PGE_2_ stimulation in both compartments (**Fig 4B**). While the soma transcriptome showed significantly increased enrichment of *translation*- and *energy*- related pathways (**Fig 4C**), these were notably downregulated in the axonal compartment. In contrast, pathways related to *receptor and ligand activity*, *inflammatory response*, *cytokine activity* and *extracellular matrix* were significantly enriched in the axonal transcriptome following PGE_2_ exposure. These findings suggest a compartmentalised inflammatory response, pointing to the axon’s distinct and active role in mediating inflammatory processes typically associated with peripheral tissue environments. In cultures of more mature neurons derived from W8 DRGs, PGE_2_ stimulation produced similar patterns of relative enrichment across subcellular domains, with increases in *translation*- and *ribosome*- related pathways in the soma and decreases in *channel activity* and *membrane-associated* processes (**Fig 5**). Unlike E16.5- derived DRG neurons, *cytokine*-related pathways were not among the most prominently enriched in the axonal transcriptomes of W8 neurons following PGE_2_ stimulation. However, W8 axons still exhibited significant reductions in *mitochondria*-related and *translation* pathways, closely mirroring the response seen in E16.5 axons. These observations suggest that mature neurons retain the capacity for axon-specific transcriptomic remodelling in response to inflammatory stimuli, and while some molecular signatures may be less prominent, the underlying regulatory themes appear largely conserved across developmental stages.

### Sensory perception, pain and axon function pathways

Building from the wider Gene Ontology (GO) analysis, we next examined specific neuronal pathways that, while not among the top-ranked terms in the initial enrichment visualisations, showed significant enrichment and are highly relevant to neuronal and nociception function. These include pathways related to *development*, *sensory perception*, *inflammation*, and *pain processing*. Their presence in the transcriptomic enrichment highlights their potential roles in the dynamic regulation of soma and axonal compartments following PGE2 stimulation.

As shown in **Supplementary** Fig 4, PGE_2_ stimulation induced a clear and consistent transcriptional response in axons, marked by significant enrichment of pathways related to *pain*, *inflammation*, and *axonal function*. In contrast, somatic transcriptomes exhibited a more generalized downregulation and reduced enrichment of these pathways. Notably, the magnitude of enrichment, particularly for pathways associated with *sensory perception* and *pain*, was more pronounced in axonal transcriptomes derived from W8 DRG neurons compared to those from E16.5-derived neurons. This contrasts with broader categories such as *structural constituent of ribosome*, *ribosomal subunit*, and *cytoplasmic translation*, which were comparably enriched in both E16.5 and W8 samples, suggesting a conserved baseline of core cellular functions. The enhanced enrichment observed in W8 axons for *sensory* and *pain-related* pathways may reflect a more differentiated and mature transcriptomic state, enabling a more specialised response to inflammatory conditions.

### Identification of functional mediators in neuronal excitability

The observed changes in subcellular axonal transcriptomes offer an opportunity to develop a data-driven pipeline for identifying molecular targets that may regulate neuronal excitability at the local, peripheral tissue level. As an initial step in identifying cellular pathways involved in DRG neuron sensitisation and functional hyperexcitability, we focused on transcripts that were axonally enriched in both E16 and W8 neurons and exhibited a significant axonal increase (>1.5-fold change and p-value < 0.05) following PGE_2_ stimulation (**Fig 6A**). Applying this stringent filtering approach yielded six candidate genes: *Ccn1*, *Gzme*, *Ngf*, *Nqo1*, *Procr*, and *Tnfrsf12a*. Notably, **Ngf** is a key mediator of peripheral pain, **CCN1**, was shown to drive pro-inflammatory programs in macrophages (Bai *et al*. 2010), and **Nqo1** has antioxidant and anti-inflammatory functions (Lee *et al*. 2021). Particularly interestingly was the strong upregulation of **Tnfrsf12a/Fn14** following PGE_2_ stimulation, a receptor implicated in nerve regeneration (Tanabe *et al*. 2003) and neuropathic pain (Huang *et al*. 2019).

**Figure 6.**
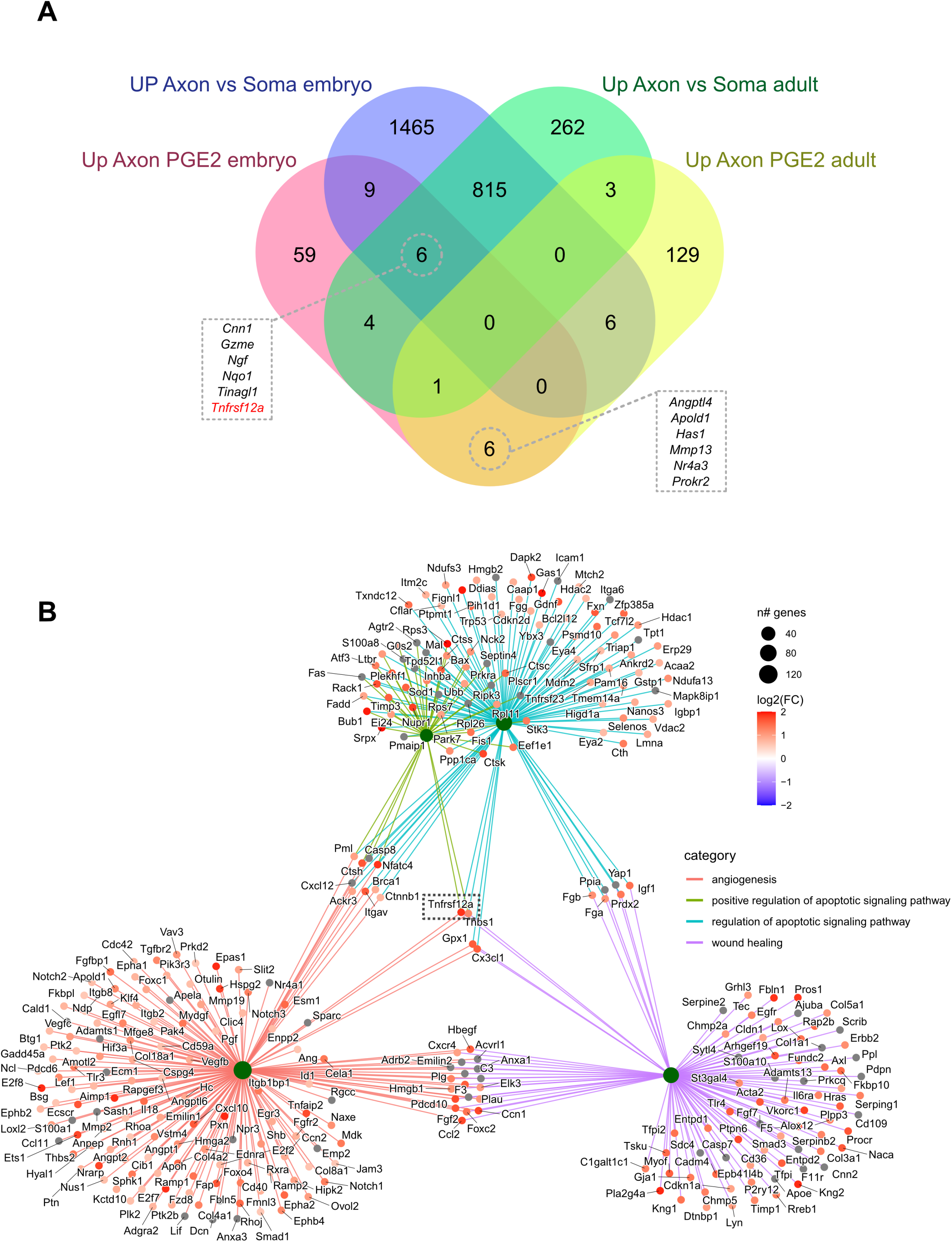
Identification of genes axonally enriched in both E16 and W8 neurons and modulated by PGE_2_. **A.** Venn diagram showing the group of genes that exhibit a significant increase (>1.5-fold change) in the axon vs soma comparison or following PGE_2_ stimulation in embryonic and adult neurons. The intersection highlights genes that are upregulated both in axon vs soma and upon PGE_2_ stimulation. **B.** Cnetplot of Gene Ontology (Biological Process) terms associated with *Tnfrsf12a* that are significantly modulated by PGE_2_ treatment in embryonic neurons. Circle color represents the log2FC value from the axon vs soma comparison; categories represent the terms, and the color of the connection lines corresponds to each term.

Our pathway analysis revealed that *Tnfrsf12a* acts in multiple functional pathways, including angiogenesis, apoptotic signalling, and wound healing, processes crucial for cellular homeostasis and survival (**Fig 6B**). It is one of only four genes (*Tnfrsf12a*, *Thbs*, *Gpx1*, and *Cx3cl*) enriched across all four highlighted pathways (**Fig 6B**). To explore the therapeutic potential of this target, we assessed how *Tnfrsf12a* regulation affects neuronal excitability using a cell-permeable siRNA probe. Target knockdown was confirmed via qPCR, with a 25% decrease in *Tnfrsf12a* expression relative to controls after 24 hours and 39% after 48 hours (fold change). The knock down of *Tnfrsf12a* expression in dissociated DRG neuron cultures (**Fig 7A**), produced a marked reduction in excitability in response to capsaicin stimulation, compared to both media and siRNA controls (**Fig 7B**). This finding highlights *Tnfrsf12a* as a promising candidate for modulating nociceptive responses.

**Figure 7.**
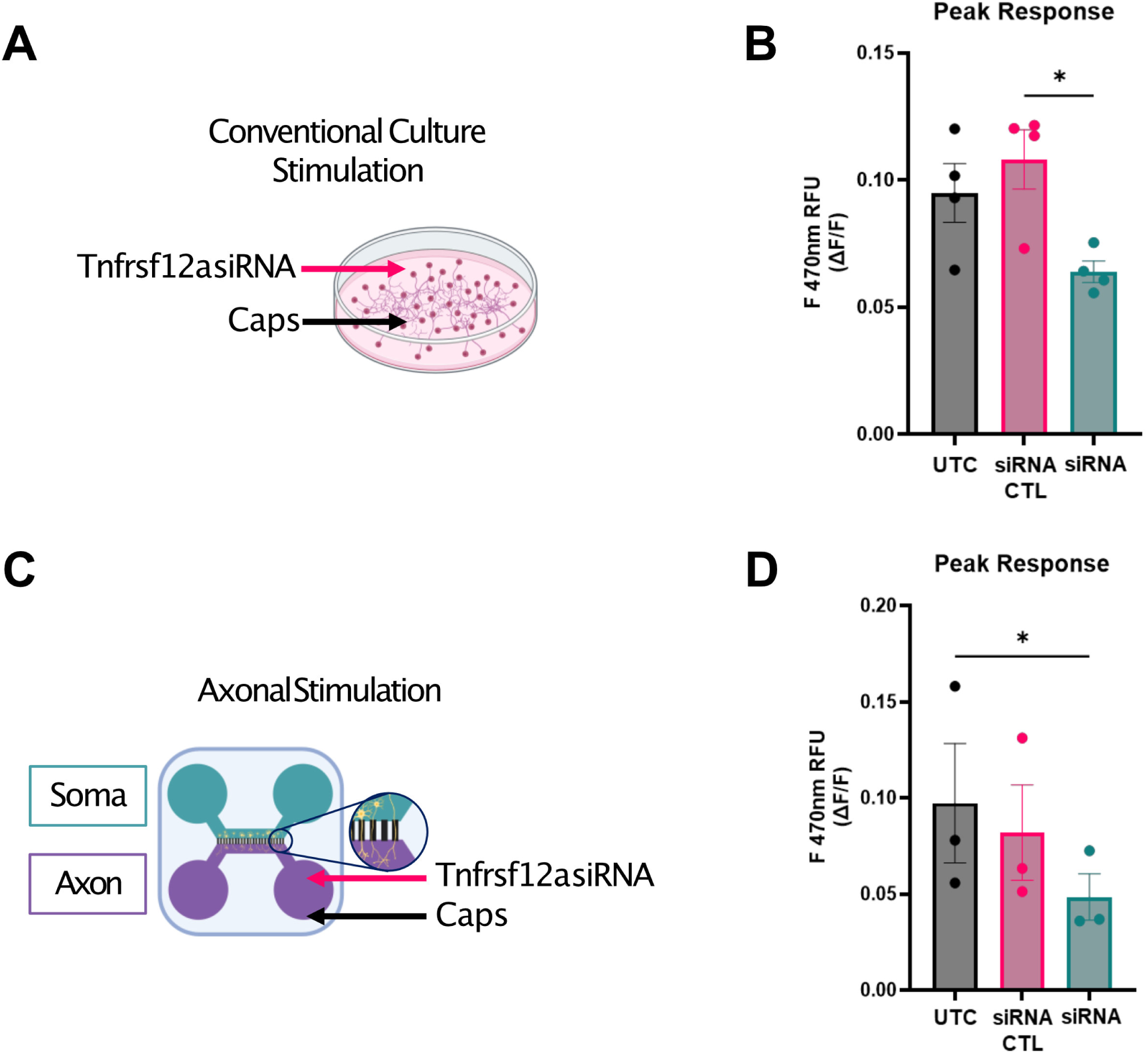
Knockdown of *Tnfrsf12a* reduces capsaicin-evoked Ca²⁺ responses in DRG neurons. **A.** Dissociated DRG sensory neurons were cultured for 7 days and incubated with 1 μM *Tnfrsf12a* siRNA or 1 μM non-targeting siRNA control for 48 hours. Untreated control (UTC) neurons received a media change only. Neurons were then stimulated with 200 nM capsaicin during Ca²⁺ imaging. **B.** Quantification of Ca²⁺ responses (ΔF/F_0_) revealed a decrease in peak responses in neurons treated with *Tnfrsf12a* siRNA compared to siRNA control, indicating diminished Ca²⁺ signalling following gene knockdown. **C.** DRG neurons were cultured in compartmentalised microfluidic chambers, and only the axonal compartment was selectively exposed to *Tnfrsf12a* siRNA or non-targeting siRNA control for 48 hours. **D.** Neurons with axons treated with *Tnfrsf12a* siRNA exhibited a significantly reduced peak Ca^2+^ response to axonal capsaicin stimulation relative to UTC. Data are presented as mean ± SEM; statistical comparisons were performed using Friedman test with Dunn’s post hoc test. *p* < 0.05 was considered statistically significant. All groups have at least *N* = 3 biological replicates, and *n* ≥ 2 technical replicates per condition.

To further investigate whether the axonal compartment is a functionally relevant site of *Tnfrsf12a* activity, we applied the siRNA probe selectively to the axonal side of microfluidic chambers (**Fig 7C**), which allowed us to assess the compartment-specific contribution of *Tnfrsf12a* to neuronal excitability. Axonal application of siRNA also resulted in a significant reduction in neuronal excitability following axon specific stimulation with capsaicin (200 nM), compared to both vehicle and control siRNA treatments (**Fig 7D**). These results underscore the functional relevance of local axonal *Tnfrsf12a* expression in modulating excitability and suggest that axonal targeting of key regulatory transcripts may provide a spatially precise strategy for therapeutic intervention in peripheral sensitisation.

## Discussion

Our findings reveal the distinct identity of axonal transcriptomes in sensory neurons, showing that a core set of transcripts essential for axonal function is preserved from embryonic (E16.5) to adult (W8) stages. This conservation supports the existence of a stable axonal transcriptome underpinning diverse functions across development. In adults, however, enriched transcripts increasingly cluster in pathways linked to *sensory processing* and *pain*, with this enrichment further amplified by PGE_2_ stimulation. These same pathways were downregulated in the soma, pointing to a previously unrecognised, compartment-specific shift in transcriptomic regulation, seen both at embryonic and adult stage. At functional level, we modelled neuronal sensitisation using prolonged PGE_2_ exposure, a paradigm more consistent with persistent pain states (Malty *et al*. 2016; Ma and St-Jacques 2018) instead of the acute treatments more commonly reported (Stucky and Lewin 1999; Southall and Vasko 2001; Kimourtzis and Raouf 2024). Compartmentalised microfluidic chambers allowed us to selectively apply PGE_2_ to peripheral axons, mimicking a local inflammatory insult. This prolonged exposure not only sensitised axons but also drove retrograde communication to the soma, priming cell bodies for heightened responses. Importantly, this platform also enabled local knockdown of *Tnfrsf12a*, an axonally enriched transcript identified in our analyses, thereby testing its functional role and providing proof-of-concept for our central hypothesis.

### DRG transcriptomes

As primary mediators of pain, DRG neurons express a wide repertoire of receptors and signalling pathways that allow them to detect environmental cues and integrate inputs from tissue-resident and immune cells during inflammation, thereby shaping pain sensitivity. Single-cell RNA sequencing has refined DRG subtype classification, demonstrating that gene expression profiles align with functional identity (Usoskin *et al*. 2015) While distinct expression patterns emerge across subtypes, many gene sets, particularly those encoding ion channels, adhesion molecules, and receptors, remain confined to specific classes (Zheng *et al*. 2019).

As axons extend into peripheral and central targets by E15.5, the identity of postmitotic DRG neurons is established, further shaped by extrinsic cues during postnatal maturation (Sharma *et al*. 2020). Yet transcriptional identity remains highly plastic, as illustrated by injury, which rapidly disrupts subtype-specific profiles but allows them to re-emerge during recovery (Nguyen *et al*. 2019). Single-nucleus RNA-seq has indeed revealed common injury-induced programs across DRG subtypes (Renthal *et al*. 2020), but axotomy also suppresses many identity-defining genes, complicating subtype resolution in injury states. Collectively, these findings highlight the remarkable plasticity of sensory neurons and underscore the importance of examining this adaptability at the subcellular, particularly axonal, level, given their complex cytoarchitecture and functional compartmentalisation.

### Subcellular DRG transcriptomes

In this context, subcellular omics approaches (Koppel and Fainzilber 2018) have significantly advanced our understanding of DRG neurons by revealing the functional relevance of compartment-specific transcriptomes. By focusing on distinct domains such as axons and neurites, these methods complement single-cell analyses and shed light on the spatiotemporal regulation of gene expression, emphasising how localised molecular processes contribute to neuronal function.

We are now beginning to understand the functional consequences of localising mRNAs and ncRNAs to distinct cellular compartments (Farias *et al*. 2019; Corradi and Baudet 2020; Paolo *et al*. 2021; Mesquita-Ribeiro *et al*. 2021; Taylor and Nikolaou 2024), as well as the impact that dysregulation of localised protein synthesis can have on the onset and development of neuronal disorders. Indeed, high-throughput sequencing studies over the last decade have identified hundreds to thousands of mRNAs that can be localised to neurites/axons across diverse neuronal types, experimental approaches and different species (Paolo *et al*. 2021). These advances were systematically reviewed in (Kügelgen and Chekulaeva 2020), where 20 datasets were re-analysed using the same bioinformatics pipeline (Wurmus *et al*. 2018). This approach enabled the identification of a core set of neurite-associated transcripts, defined as those detected in at least 16 out of the 20 datasets.

Applying this list of core transcripts to our own DRG RNA-seq dataset from axons/neurites, we were able to demonstrate a clear enrichment of this core neurite transcriptome signature. These included RNAs encoding ribosomal proteins (Rps2, Rps12, Rps18, Fau, Rplp1), as well as RNAs with a variety of localised roles, such as Cox6a1 (mitochondrial function), Usf2 (Ca^2+^ responsive transcription), Nes (outgrowth formation), Anp32b (nuclear protein), Rhoc (plasma membrane), plus Ybx1 and Eif3f, known for their roles in RNA binding and translation machinery respectively.

One of the key features now broadly documented in neurite and axon transcriptomes is the presence of mRNAs coding for proteins involved in nuclear functions, particularly transcription factors (TFs) and components of the nuclear transport machinery. This observation underscores the role of axonally translated proteins in retrograde signalling, where they can influence gene expression profiles and thereby modulate axonal function (Cox *et al*. 2008; Ben-Yaakov *et al*. 2012; Perry and Fainzilber 2014; Rangaraju *et al*. 2017; Terenzio *et al*. 2018; Lucci *et al*. 2020). For example, in both our E16.5 and W8 axonal transcriptomes, we show high expression of TFs Atf4 and Creb1 (**Supplementary Tables 2-3**), which have been previously associated with local translation and pain pathways (Cox *et al*. 2008; Dong *et al*. 2011; Baleriola *et al*. 2014; Cagnetta *et al*. 2019).

Furthermore, previous studies showed how elevated expression of Atf4 RNA in a model of retinal cell degeneration was associated with increased expression of inflammatory and nociceptive mediators IL-1B, IL-6 and TNF-a, in a pathway mediated by eIF2-a (Rana *et al*. 2014).

Although no differentially expressed genes (DEGs) were identified after multiple testing correction, GSEA provided a robust alternative by revealing coordinated, pathway-level changes across the transcriptome, independent of individual gene-level significance. These analyses reinforce the now well-established presence of nuclear protein-encoding transcripts within axons. Notably, our axonal DRG datasets show strong enrichment for transcripts associated with the translational and ribosomal machinery, as well as genes involved in mitochondrial function and energy production (**Figs 2-3**). This finding highlights the high translational and metabolic demands of actively growing and regenerating axons, demands that are a common characteristic of DRG neurons at both embryonic and adult stages. At mechanistic level, the striking axonal enrichment of transcripts encoding *ribosomal proteins* (RPs), which were typically thought to be assembled solely in the nucleus, reflects a shift in our understanding of axonal translation. Indeed, recent insights suggest that axonally synthesized RPs are directly associated with functional ribosomes in the axon, independent of the nucleolus (Shigeoka *et al*. 2019; Nagano and Araki 2021). This emphasises the likelihood that axonal transcriptomes are translated by locally assembled and specialised ribosomes, highlighting the axon’s capacity for independent protein synthesis, and the potential for axon-specific regulatory mechanisms. In this regard, previous work by the Holt group (Shigeoka *et al*. 2019) uncovered crucial functions of the locally synthesised ribosomal protein Rps4x in axonal mRNA translation and in axon branching in vivo. Indeed, their previous work (Shigeoka *et al*. 2016) had showed how the number of axonally translated mRNAs is highest during branching stages, when synapses are being formed. This important emerging role of Rps4x in axonal protein synthesis is reinforced by our findings in DRG neurons, where it ranks as one of the most highly enriched transcripts in axons.

In addition to RPs, our axonal transcriptomes show significant enrichment of transcripts encoding mitochondrial proteins, a trend also observed in previous axonal datasets. This pattern holds across our comparison of E16.5 and W8 DRG axonal transcriptomes, suggesting that these proteins may play crucial roles in axonal function. Collectively, these findings underscore the metabolically demanding and translationally intensive processes that are essential for axonal function and growth.

### Dynamic changes in DRG transcriptomes after injury and inflammatory process

The expression patterns of mRNAs localised to the axon has been shown to correlate with the probability of local translation (Shigeoka *et al*. 2016; Zappulo *et al*. 2017), and the existence of a distinct axonal transcriptome subject to changes in neuronal sensitisation would offer the possibility of interventions delivered specifically to peripheral tissues (i.e. OA knee). In this context, the characterisation of the axonal transcriptome following PGE_2_-driven sensitisation enables the search for locally relevant transcripts. However, despite the recognition that local translation is a crucial requirement for axonal growth/regeneration (Verma *et al*. 2005), and the fact that studies such as those involving rat paw injections of protein synthesis inhibitors like rapamycin, cordycepin, and anisomycin have demonstrated the importance of localised protein translation in the development of hyperalgesia in vivo (Jiménez-Díaz *et al*. 2008; Bogen *et al*. 2012; Ferrari *et al*. 2013), the search for novel pain targets has yet to prioritise those transcripts that are axonally localised.

Our findings show how transcripts identified as signature axon/neurite markers are decreased in the axon, while their comparatively low levels in the soma transcriptome show a general increase (**Figs 4-5**). Indeed, despite how injury models like CFA-triggered inflammatory pain do not induce a transcriptional state comparable to that observed after axotomy (Renthal *et al*. 2020), it has been suggested that axonal injury may reactivate embryonic developmental programs to promote regeneration (Harel and Strittmatter 2006; Poplawski *et al*. 2020). This demonstrates differential responses to transcriptional reprogramming across specific neuronal domains, a phenomenon evident not only at the level of axonal markers, but also, importantly, at the global expression level. For instance, while the soma transcriptome shows enrichment of *translation* and *ribosomal*- associated pathways following PGE_2_ stimulation, axonal changes exhibit a contrasting pattern, with significant decreases in pathways linked with *respiration* and *translation*. Notably, *cytokine activity*, *inflammatory response*, and *receptor activity* are all upregulated at the axonal level, highlighting the presence of locally triggered responses that are distinct from those occurring in the soma.

Following PGE_2_ stimulation, axonal transcriptomes show increased enrichment of transcripts associated with *neuronal perception* and *axon*-related processes, a pattern not observed in soma transcriptomes (**Fig 6**). These distinct axonal changes may offer a more translationally relevant model for identifying novel pain therapy targets. Given the axon’s direct role in sensory processing and pain transmission, focusing on transcriptomic alterations within this compartment could uncover molecular pathways more closely linked to the onset and persistence of pain.

### Modulating Axonal Transcripts to Control neuronal responses to noxious stimuli: Tnfrsf12a as a Target

To identify proof-of-concept candidates demonstrating the role of axonal transcripts in neuronal responses to noxious stimuli, we implemented a stringent selection pipeline, focusing exclusively on transcripts both enriched in axons and upregulated following PGE_2_ stimulation. Among these, we prioritized *Tnfrsf12a* (also known as *Fn14* or *TWEAKR*), the receptor for *Tnfsf12* (TWEAK), a TNF superfamily ligand expressed in both membrane-bound and soluble forms. Unlike other TNF ligands, *Tnfsf12* is broadly expressed, upregulated in multiple injury models, and involved in a range of cellular processes including proliferation, migration, survival, differentiation, and cell death (Winkles 2008).

Tnfrsf12a is the smallest member of the TNF-receptor superfamily and has been shown to increase after injury, functioning as part of a multifunctional pro-inflammatory and pro-angiogenic pathway (Winkles 2008). In DRG neurons, early studies found that *Tnfrsf12a* is induced in axotomized neurons and promotes neurite outgrowth, suggesting a potential role in nerve regeneration (Tanabe *et al*. 2003). More recently, *Tnfrsf12* expression at both mRNA and protein levels was shown to increase in the ipsilateral DRG following spinal nerve ligation, with intrathecal administration of a Tnfrsf12 protein inhibitor or DRG knockdown alleviating neuropathic pain (Huang *et al*. 2019). We reveal that *Tnfrsf12a* is not only regulated by inflammatory signals, leading to increased mRNA expression, but also that its function extends beyond previously described roles. Through subcellular transcriptomics, we uncovered a distinct local role for *Tnfrsf12a* translation at the axonal level, where inflammatory signals may drive sensory hyperexcitability in pain conditions.

Our data align and support previous research on neuronal injury (Rishal and Fainzilber 2014; Prior *et al*. 2017; Pizio *et al*. 2022) by demonstrating that axonal transcriptomes retain a conserved identity from embryonic to adult stages but are also capable of dynamically responding to inflammatory stimuli such as PGE_2_. Tnfrsf12a (FN14) may play a dual role in both regenerative and maladaptive nociceptive processes, similar to other TNF superfamily members involved in neuroinflammatory responses. Moreover, the shift in transcriptional regulation between soma and axon following PGE_2_ stimulation suggests a compartmentalised response to inflammatory cues, reinforcing the concept that axonal signaling mechanisms are integral to neuronal adaptation and disease progression.

These findings provide further evidence that subcellular transcriptomics can uncover novel aspects of injury signaling and highlight the importance of local translation in sensory neuron plasticity.

Modulating axonal plasticity and dynamics to control the firing potential that triggers pain in chronic conditions presents a promising therapeutic avenue for managing persistent pain. Targeting key axonal transcripts that respond to inflammatory cues and influence sensory hyperexcitability could provide new strategies for intervention. Crucially, prioritising the development of drugs that act specifically on peripheral nerve terminals offers a significant advantage, allowing for effective pain relief while minimizing unwanted effects on central nervous system function. By leveraging insights from subcellular transcriptomics, future therapies may achieve greater precision in disrupting pathological pain signalling at its source.

## Acknowledgements and Funding

We thank the Ted Price laboratory (University of Texas at Dallas) for their support with analytical pipelines. This work was funded by the Pain Centre Versus Arthritis (grant reference 20777), a Versus Arthritis PhD Scholarship (grant reference 21586; C.dM, grant holder; A.A-T, awardee), and the University of Nottingham BBSRC Doctoral Training Programme (RP, awardee).

## Supplementary Materials

**Supplementary Figure 1.**
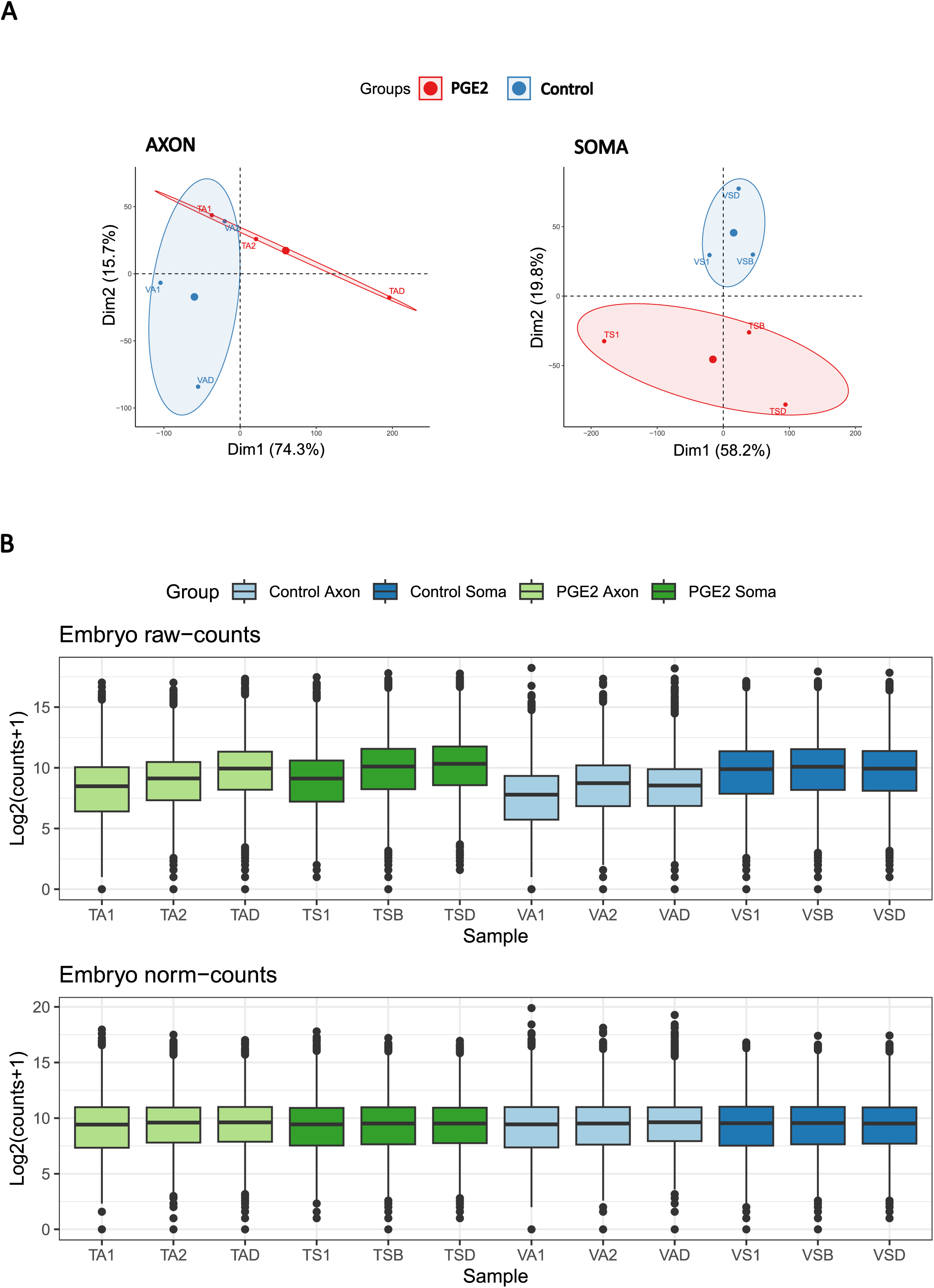
PCA plots and boxplots of raw and normalized RNA-seq data from embryo datasets. **A.** PCA plot of Axon and Soma samples using raw counts. **B.** Boxplots of samples before (raw counts) and after normalization (normalized counts).

**Supplementary Figure 2.**
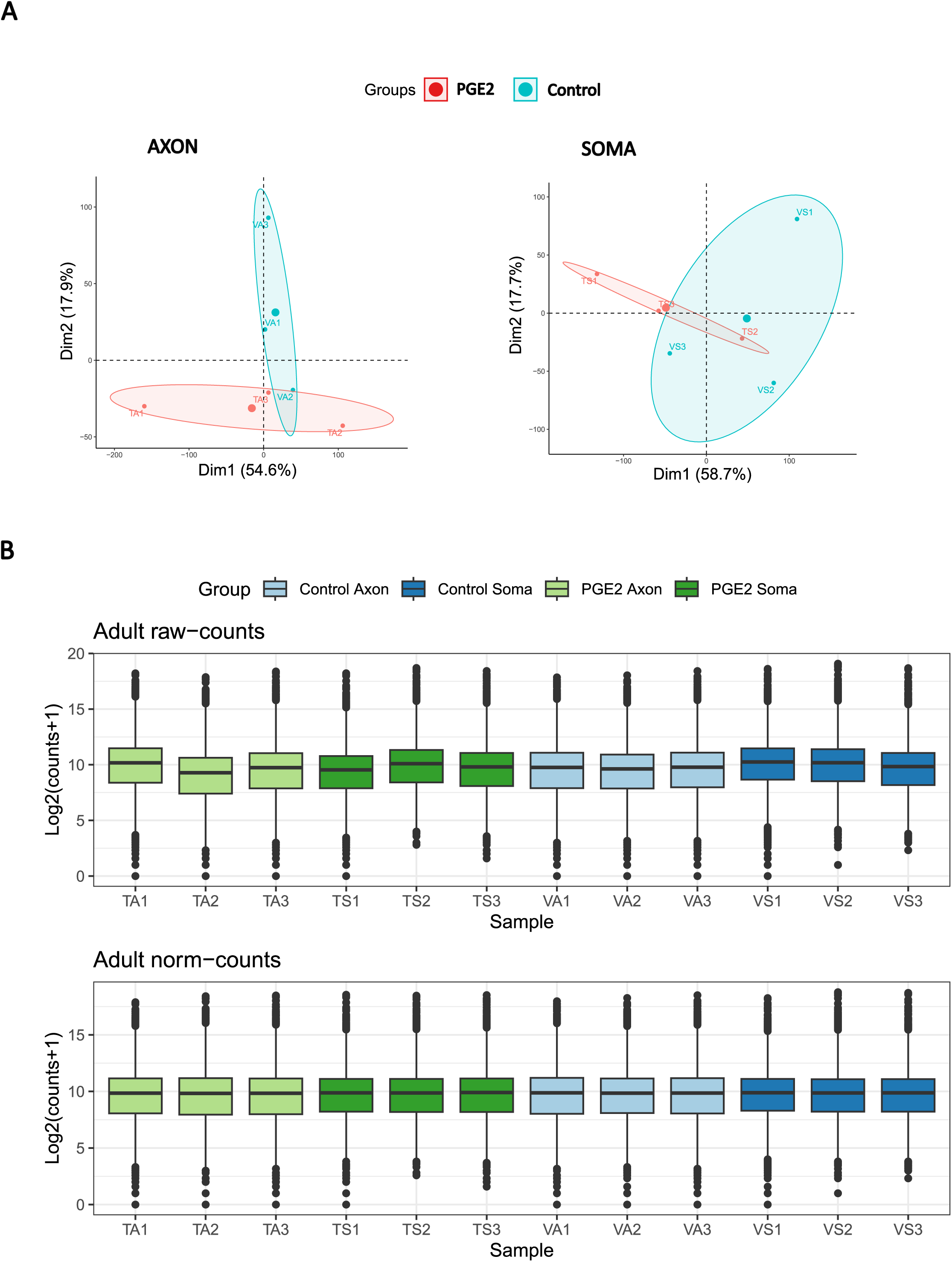
PCA plots and boxplots of raw and normalized RNA-seq data from adult datasets. **A.** PCA plot of Axon and Soma samples using raw counts. **B.** Boxplots of samples before (raw counts) and after normalization (normalized counts).

**Supplementary Figure 3.**
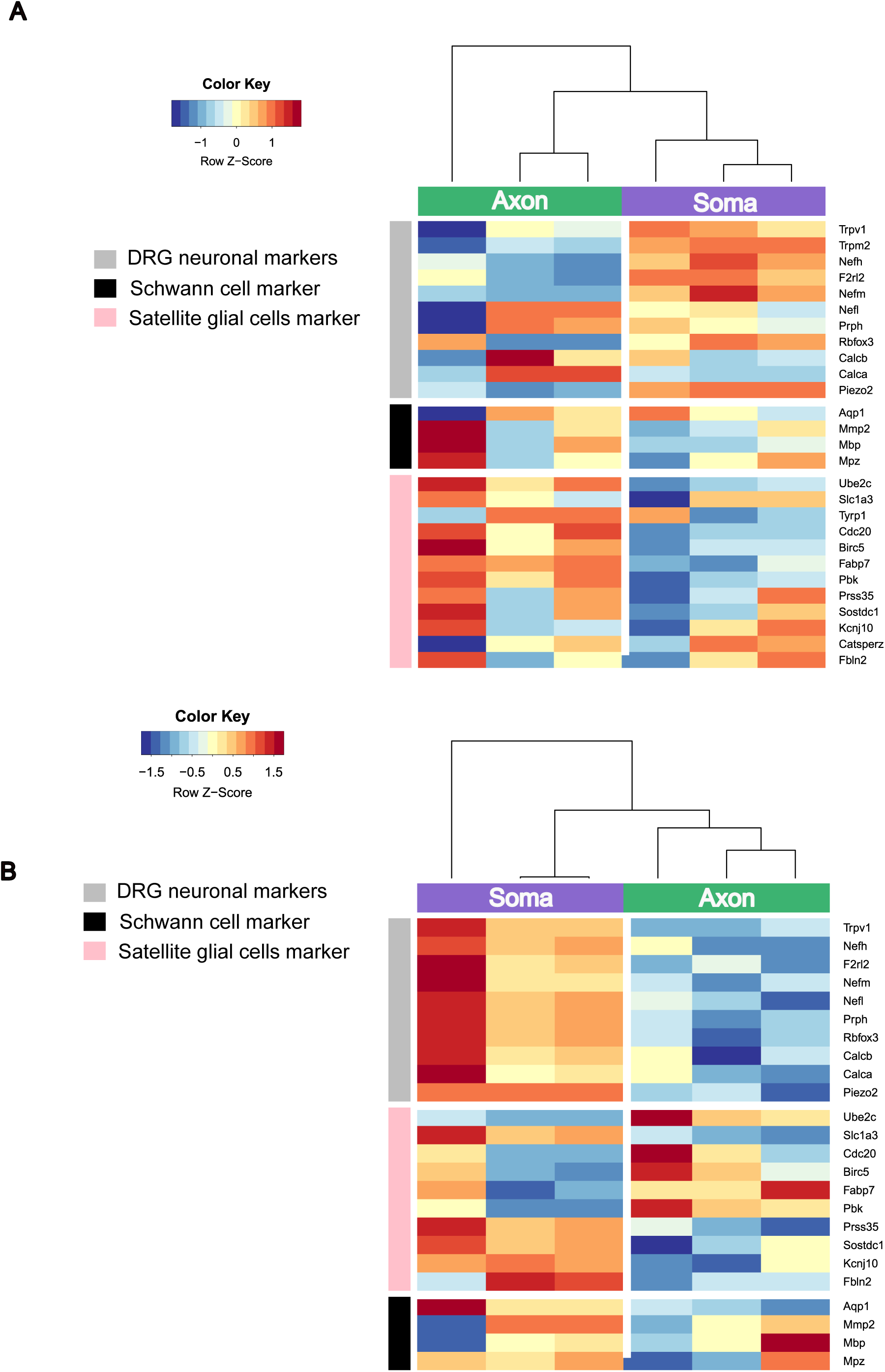
Heatmap of different cell type markers for (A) embryo and (B) adult datasets. Gene expression of 4 markers for Schwann cells, 12 markers for satellite glial cells and 11 markers for DRG neurons are shown as Z-score of log2 normalized counts (hierarchical clustering distance measured by Euclidean and Ward clustering algorithms).

**Supplementary Figure 4.**
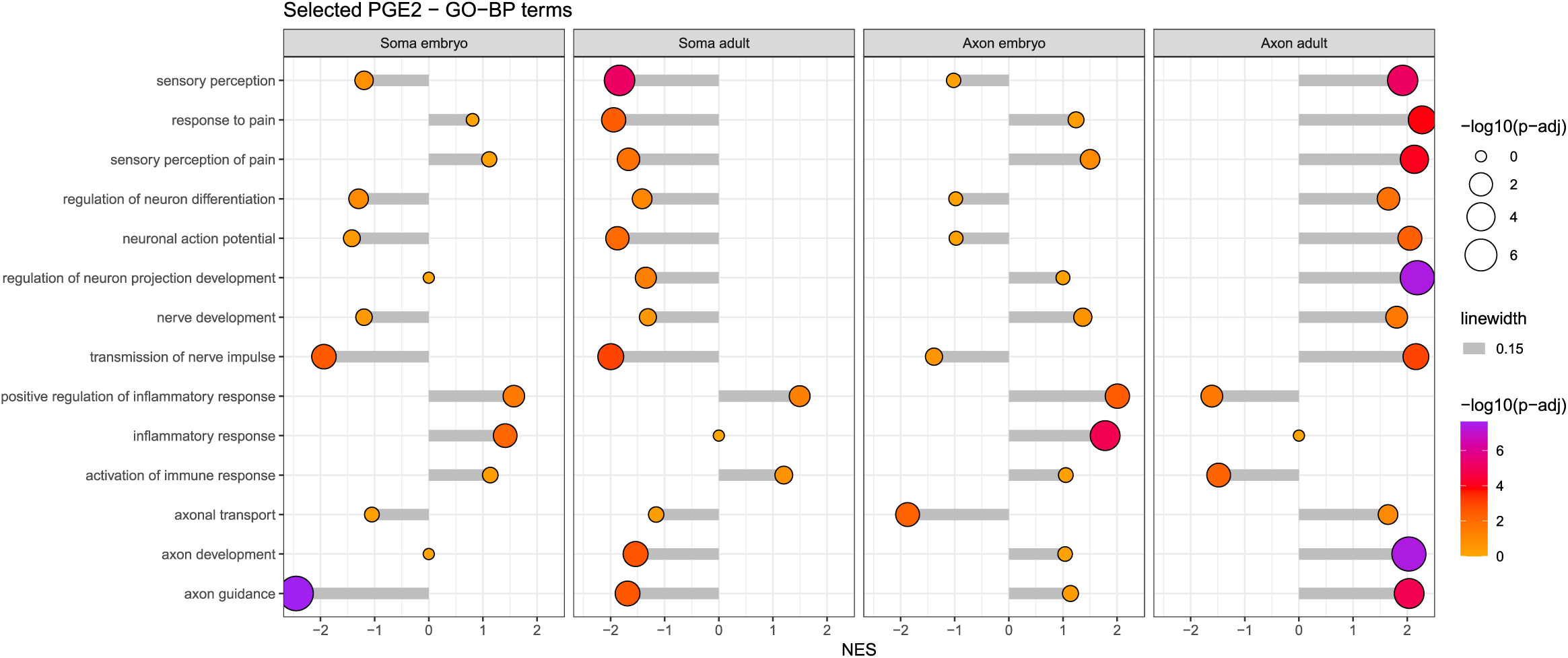
Lollipop plot of selected GSEA categories related to pain in embryonic and adult datasets treated with PGE_2_. Selected GO-BP (related to development, sensory perception, inflammation, and pain processing) terms enriched in PGE2-regulated transcriptomes are shown for soma (embryo and adult) and axon (embryo and adult) datasets. The x-axis represents the normalized enrichment score (NES), where positive values indicate upregulation and negative values indicate downregulation of the corresponding process. Circle size and color is proportional to the statistical significance (−log10 adjusted p-value).

**Supplementary Table 1: Statistics of sample filtering and mapping to the mouse genome.**

**Supplementary Table 2: Differentially expressed genes (DEGs) analysis in embryonic samples.**

**Supplementary Table 3: Differentially expressed genes (DEGs) analysis in adult samples.**

## References

Bai T., Chen C.-C., Lau L. F. (2010) Matricellular Protein CCN1 Activates a Proinflammatory Genetic Program in Murine Macrophages. J. Immunol. 184, 3223–3232.

Baleriola J., Walker C. A., Jean Y. Y., Crary J. F., Troy C. M., Nagy P. L., Hengst U. (2014) Axonally synthesized ATF4 transmits a neurodegenerative signal across brain regions. Cell 158, 1159– 1172.

Banchet G. S. von, Kiehl M., Schaible H. (2005) Acute and long-term effects of IL-6 on cultured dorsal root ganglion neurones from adult rat. J. Neurochem. 94, 238–248.

Basbaum A. I., Bautista D. M., Scherrer G., Julius D. (2009) Cellular and molecular mechanisms of pain. Cell 139, 267–284.

Ben-Yaakov K., Dagan S. Y., Segal-Ruder Y., Shalem O., Vuppalanchi D., Willis D. E., Yudin D., et al. (2012) Axonal transcription factors signal retrogradely in lesioned peripheral nerve. The EMBO Journal 31, 1350–1363.

Bogen O., Alessandri-Haber N., Chu C., Gear R. W., Levine J. D. (2012) Generation of a pain memory in the primary afferent nociceptor triggered by PKCε activation of CPEB. J Neurosci 32, 2018– 2026.

Cagnetta R., Wong H. H.-W., Frese C. K., Mallucci G. R., Krijgsveld J., Holt C. E. (2019) Noncanonical Modulation of the eIF2 Pathway Controls an Increase in Local Translation during Neural Wiring. Mol Cell 73, 474–489.e5.

Chaban V. V. (2010) Peripheral sensitization of sensory neurons. Ethn. Dis. 20, S1–3–6.

Chakrabarti S., Hore Z., Pattison L. A., Lalnunhlimi S., Bhebhe C. N., Callejo G., Bulmer D. C., Taams L. S., Denk F., Smith E. St. J. (2020) Sensitization of knee-innervating sensory neurons by tumor necrosis factor-α-activated fibroblast-like synoviocytes: an in vitro, coculture model of inflammatory pain. Pain 161, 2129–2141.

Clark P., Rowland S. E., Denis D., Mathieu M.-C., Stocco R., Poirier H., Burch J., et al. (2008) MF498 [N -{[4-(5,9-Diethoxy-6-oxo-6,8-dihydro-7 H -pyrrolo[3,4-g]quinolin-7-yl)-3- methylbenzyl]sulfonyl}-2-(2-methoxyphenyl)acetamide], a Selective E Prostanoid Receptor 4 Antagonist, Relieves Joint Inflammation and Pain in Rodent Models of Rheumatoid and Osteoarthritis. J. Pharmacol. Exp. Ther. 325, 425–434.

Corradi E., Baudet M.-L. (2020) In the Right Place at the Right Time: miRNAs as Key Regulators in Developing Axons. Int J Mol Sci 21, 8726–22.

Cox L. J., Hengst U., Gurskaya N. G., Lukyanov K. A., Jaffrey S. R. (2008) Intra-axonal translation and retrograde trafficking of CREB promotes neuronal survival. Nat Cell Biol 10, 149–159.

Deglincerti A., Jaffrey S. R. (2012) Insights into the roles of local translation from the axonal transcriptome. Open Biol 2, 120079–120079.

Dong L., Guarino B. B., Jordan-Sciutto K. L., Winkelstein B. A. (2011) Activating transcription factor 4, a mediator of the integrated stress response, is increased in the dorsal root ganglia following painful facet joint distraction. Neuroscience 193, 377–386.

Farias J., Sotelo J. R., Sotelo-Silveira J. (2019) Toward Axonal System Biology: Genome Wide Views of Local mRNA Translation. Proteomics 19, e1900054.

Ferrari L. F., Bogen O., Chu C., Levine J. D. (2013) Peripheral administration of translation inhibitors reverses increased hyperalgesia in a model of chronic pain in the rat. J Pain 14, 731–738.

Finnerup N. B., Kuner R., Jensen T. S. (2021) Neuropathic Pain: From Mechanisms to Treatment. Physiol. Rev. 101, 259–301.

Garcez P. P., Guillemot F., Dajas-Bailador F. (2016) MicroRNA Technologies. Neuromethods, 59–71.

Gardiner N. J., Freeman O. J. (2016) Chapter Five Can Diabetic Neuropathy Be Modeled In Vitro? Int. Rev. Neurobiol. 127, 53–87.

Harel N. Y., Strittmatter S. M. (2006) Can regenerating axons recapitulate developmental guidance during recovery from spinal cord injury? Nat Rev Neurosci 7, 603–616.

Huang L.-N., Zou Y., Wu S.-G., Zhang H.-H., Mao Q.-X., Li J.-B., Tao Y.-X. (2019) Fn14 Participates in Neuropathic Pain Through NF-κB Pathway in Primary Sensory Neurons. Mol Neurobiol 56, 7085– 7096.

Jiménez-Díaz L., Géranton S. M., Passmore G. M., Leith J. L., Fisher A. S., Berliocchi L., Sivasubramaniam A. K., Sheasby A., Lumb B. M., Hunt S. P. (2008) Local translation in primary afferent fibers regulates nociception. PLoS ONE 3, e1961.

Khoutorsky A., Price T. J. (2018) Translational Control Mechanisms in Persistent Pain. Trends Neurosci. 41, 100–114.

Kimourtzis G., Raouf R. (2024) A microfluidic model of the first sensory synapse for analgesic target discovery. Mol. Pain 20, 17448069241293286.

Koppel I., Fainzilber M. (2018) Omics approaches for subcellular translation studies. *Mol*. Omics 14, 380–388.

Kügelgen N., Chekulaeva M. (2020) Conservation of a core neurite transcriptome across neuronal types and species. Wiley Interdiscip Rev Rna 11, e1590.

Latremoliere A., Woolf C. J. (2009) Central Sensitization: A Generator of Pain Hypersensitivity by Central Neural Plasticity. J. Pain 10, 895–926.

Lee W.-S., Ham W., Kim J. (2021) Roles of NAD(P)H:quinone Oxidoreductase 1 in Diverse Diseases. Life 11, 1301.

Li X., Jin D. S., Eadara S., Caterina M. J., Meffert M. K. (2023) Regulation by noncoding RNAs of local translation, injury responses, and pain in the peripheral nervous system. Neurobiology Pain 13, 100119.

Liang Z., Hore Z., Harley P., Stanley F. U., Michrowska A., Dahiya M., Russa F. L., Jager S. E., Villa-Hernandez S., Denk F. (2020) A transcriptional toolbox for exploring peripheral neuroimmune interactions. Pain 161, 2089–2106.

Lucci C., Mesquita-Ribeiro R., Rathbone A., Dajas-Bailador F. (2020) Spatiotemporal regulation of GSK3β levels by miRNA-26a controls axon development in cortical neurons. Development 147, dev180232-27.

Ma W., St-Jacques B. (2018) Signalling transduction events involved in agonist-induced PGE2/EP4 receptor externalization in cultured rat dorsal root ganglion neurons. Eur. J. Pain 22, 845–861.

Malty R. H., Hudmon A., Fehrenbacher J. C., Vasko M. R. (2016) Long-term exposure to PGE2 causes homologous desensitization of receptor-mediated activation of protein kinase A. J. Neuroinflammation 13, 181.

Melemedjian O. K., Khoutorsky A. (2015) Translational control of chronic pain. Prog Mol Biol Transl Sci 131, 185–213.

Mesquita-Ribeiro R., Fort R. S., Rathbone A., Farias J., Lucci C., James V., Sotelo-Silveira J., Duhagon M. A., Dajas-Bailador F. (2021) Distinct small non-coding RNA landscape in the axons and released extracellular vesicles of developing primary cortical neurons and the axoplasm of adult nerves. RNA Biology, 1–24.

Nagano S., Araki T. (2021) Axonal Transport and Local Translation of mRNA in Neurodegenerative Diseases. Front. Mol. Neurosci. 14, 697973.

Nguyen M. Q., Pichon C. E. L., Ryba N. (2019) Stereotyped transcriptomic transformation of somatosensory neurons in response to injury. eLife 8, e49679.

Obara I., Géranton S. M., Hunt S. P. (2012) Axonal protein synthesis: a potential target for pain relief? Curr Opin Pharmacol 12, 42–48.

Paolo A. D., Garat J., Eastman G., Farias J., Dajas-Bailador F., Smircich P., Sotelo-Silveira J. R. (2021) Functional Genomics of Axons and Synapses to Understand Neurodegenerative Diseases. Front. Cell. Neurosci. 15, 686722.

Peeraer E., Lutsenborg A. V., Verheyen A., Jongh R. D., Nuydens R., Meert T. F. (2011) Pharmacological evaluation of rat dorsal root ganglion neurons as an in vitro model for diabetic neuropathy. J. Pain Res. 4, 55–65.

Perry R. B., Fainzilber M. (2014) Local translation in neuronal processes—in vivo tests of a “heretical hypothesis.” Dev. Neurobiol. 74, 210–217.

Pizio A. D., Marvaldi L., Birling M.-C., Okladnikov N., Dupuis L., Fainzilber M., Rishal I. (2022) A conditional null allele of Dync1h1 enables targeted analyses of dynein roles in neuronal length sensing. J. Cell Sci. 136, jcs260220.

Poplawski G. H. D., Kawaguchi R., Niekerk E. V., Lu P., Mehta N., Canete P., Lie R., et al. (2020) Injured adult neurons regress to an embryonic transcriptional growth state. Nature 581, 77–82.

Price T. J., Géranton S. M. (2009) Translating nociceptor sensitivity: the role of axonal protein synthesis in nociceptor physiology. Eur J Neurosci 29, 2253–2263.

Price T. J., Inyang K. E. (2015) Commonalities between pain and memory mechanisms and their meaning for understanding chronic pain. Prog Mol Biol Transl Sci 131, 409–434.

Price T. J., Rashid M. H., Millecamps M., Sanoja R., Entrena J. M., Cervero F. (2007) Decreased Nociceptive Sensitization in Mice Lacking the Fragile X Mental Retardation Protein: Role of mGluR1/5 and mTOR. J. Neurosci. 27, 13958–13967.

Prior R., Helleputte L. V., Benoy V., Bosch L. V. D. (2017) Defective axonal transport: A common pathological mechanism in inherited and acquired peripheral neuropathies. Neurobiol. Dis. 105, 300–320.

Rana T., Shinde V. M., Starr C. R., Kruglov A. A., Boitet E. R., Kotla P., Zolotukhin S., Gross A. K., Gorbatyuk M. S. (2014) An activated unfolded protein response promotes retinal degeneration and triggers an inflammatory response in the mouse retina. Cell Death Dis. 5, e1578–e1578.

Rangaraju V., Dieck S. tom, Schuman E. M. (2017) Local translation in neuronal compartments: how local is local? EMBO Rep. 18, 693–711.

Ray P., Torck A., Quigley L., Wangzhou A., Neiman M., Rao C., Lam T., et al. (2018) Comparative transcriptome profiling of the human and mouse dorsal root ganglia: an RNA-seq-based resource for pain and sensory neuroscience research. PAIN® 159, 1325–1345.

Renthal W., Tochitsky I., Yang L., Cheng Y.-C., Li E., Kawaguchi R., Geschwind D. H., Woolf C. J. (2020) Transcriptional Reprogramming of Distinct Peripheral Sensory Neuron Subtypes after Axonal Injury. Neuron 108, 128–144.e9.

Ricciotti E., FitzGerald G. A. (2011) Prostaglandins and Inflammation. *Arter., Thromb., Vasc*. Biol. 31, 986–1000.

Rishal I., Fainzilber M. (2014) Axon-soma communication in neuronal injury. Nat Rev Neurosci 15, 32–42.

Rush A. M., Waxman S. G. (2004) PGE2 increases the tetrodotoxin-resistant Nav1.9 sodium current in mouse DRG neurons via G-proteins. Brain Res. 1023, 264–271.

Sahoo P. K., Smith D. S., Perrone-Bizzozero N., Twiss J. L. (2018) Axonal mRNA transport and translation at a glance. J Cell Sci 131, jcs196808.

Schmittgen T. D., Livak K. J. (2008) Analyzing real-time PCR data by the comparative C(T) method. Nat Protoc 3, 1101–1108.

Sharma N., Flaherty K., Lezgiyeva K., Wagner D. E., Klein A. M., Ginty D. D. (2020) The emergence of transcriptional identity in somatosensory neurons. Nature 577, 392–398.

Shigeoka T., Jung H., Jung J., Turner-Bridger B., Ohk J., Lin J. Q., Amieux P. S., Holt C. E. (2016) Dynamic Axonal Translation in Developing and Mature Visual Circuits. Cell 166, 181–192.

Shigeoka T., Koppers M., Wong H. H.-W., Lin J. Q., Cagnetta R., Dwivedy A., Nascimento J. de F., et al. (2019) On-Site Ribosome Remodeling by Locally Synthesized Ribosomal Proteins in Axons. Cell Rep 29, 3605–3619.e10.

Southall M. D., Vasko M. R. (2001) Prostaglandin Receptor Subtypes, EP3C and EP4, Mediate the Prostaglandin E2-induced cAMP Production and Sensitization of Sensory Neurons*. J. Biol. Chem. 276, 16083–16091.

Stucky C. L., Lewin G. R. (1999) Isolectin B(4)-positive and -negative nociceptors are functionally distinct. J Neurosci 19, 6497–6505.

Subramanian A., Tamayo P., Mootha V. K., Mukherjee S., Ebert B. L., Gillette M. A., Paulovich A., et al. (2005) Gene set enrichment analysis: A knowledge-based approach for interpreting genome-wide expression profiles. Proc. Natl. Acad. Sci. 102, 15545–15550.

Tanabe K., Bonilla I., Winkles J. A., Strittmatter S. M. (2003) Fibroblast growth factor-inducible-14 is induced in axotomized neurons and promotes neurite outgrowth. J Neurosci 23, 9675–9686.

Taylor R., Nikolaou N. (2024) RNA in axons, dendrites, synapses and beyond. Front. Mol. Neurosci. 17, 1397378.

Terenzio M., Koley S., Samra N., Rishal I., Zhao Q., Sahoo P. K., Urisman A., et al. (2018) Locally translated mTOR controls axonal local translation in nerve injury. Science 359, 1416–1421.

Unsain N., Heard K. N., Higgins J. M., Barker P. A. (2014) Production and Isolation of Axons from Sensory Neurons for Biochemical Analysis Using Porous Filters. J. Vis. Exp. 89.

Usoskin D., Furlan A., Islam S., Abdo H., Lönnerberg P., Lou D., Hjerling-Leffler J., et al. (2015) Unbiased classification of sensory neuron types by large-scale single-cell RNA sequencing. Nat. Neurosci. 18, 145–153.

Verma P., Chierzi S., Codd A. M., Campbell D. S., Meyer R. L., Holt C. E., Fawcett J. W. (2005) Axonal protein synthesis and degradation are necessary for efficient growth cone regeneration. J Neurosci 25, 331–342.

Wangzhou A., McIlvried L. A., Paige C., Barragan-Iglesias P., Shiers S., Ahmad A., Guzman C. A., et al. (2020) Pharmacological target-focused transcriptomic analysis of native vs cultured human and mouse dorsal root ganglia. Pain 161, 1497–1517.

Winkles J. A. (2008) The TWEAK–Fn14 cytokine–receptor axis: discovery, biology and therapeutic targeting. Nat Rev Drug Discov 7, 411–425.

Wu T., Hu E., Xu S., Chen M., Guo P., Dai Z., Feng T., et al. (2021) clusterProfiler 4.0: A universal enrichment tool for interpreting omics data. Innov. 2, 100141.

Wurmus R., Uyar B., Osberg B., Franke V., Gosdschan A., Wreczycka K., Ronen J., Akalin A. (2018) PiGx: reproducible genomics analysis pipelines with GNU Guix. GigaScience 7, giy123.

Zappulo A., Bruck D. van den, Mattioli C. C., Franke V., Imami K., McShane E., Moreno-Estelles M., et al. (2017) RNA localization is a key determinant of neurite-enriched proteome. Nature Communications 8, 583–13.

Zeisel A., Hochgerner H., Lönnerberg P., Johnsson A., Memic F., Zwan J. van der, Häring M., et al. (2018) Molecular Architecture of the Mouse Nervous System. Cell 174, 999–1014.e22.

Zheng Y., Liu P., Bai L., Trimmer J. S., Bean B. P., Ginty D. D. (2019) Deep Sequencing of Somatosensory Neurons Reveals Molecular Determinants of Intrinsic Physiological Properties. Neuron 103, 598–616.e7.

